# Predicting and Interpreting Protein Developability via Transfer of Convolutional Sequence Representation

**DOI:** 10.1101/2022.11.21.517400

**Authors:** Alexander W. Golinski, Zachary D. Schmitz, Gregory H. Nielsen, Bryce Johnson, Diya Saha, Sandhya Appiah, Benjamin J. Hackel, Stefano Martiniani

**Affiliations:** Department of Chemical Engineering and Materials Science, University of Minnesota, Minneapolis, MN 55455; Center for Soft Matter Research, Department of Physics, New York University, New York, NY 10003; Simons Center for Computational Physical Chemistry, Departments of Chemistry, New York University, New York, NY 10003; Courant Institute of Mathematical Sciences, New York University, New York, NY 10003

## Abstract

Engineered proteins have emerged as novel diagnostics, therapeutics, and catalysts. Often, poor protein developability – quantified by expression, solubility, and stability – hinders utility. The ability to predict protein developability from amino acid sequence would reduce the experimental burden when selecting candidates. Recent advances in screening technologies enabled a high-throughput developability dataset for 10^5^ of 10^20^ possible variants of protein ligand scaffold Gp2. In this work, we evaluate the ability of neural networks to learn a developability representation from a high-throughput dataset and transfer this knowledge to predict recombinant expression beyond observed sequences. The model convolves learned amino acid properties to predict expression levels 44% closer to the experimental variance compared to a non-embedded control. Analysis of learned amino acid embeddings highlights the uniqueness of cysteine, the importance of hydrophobicity and charge, and the unimportance of aromaticity, when aiming to improve the developability of small proteins. We identify clusters of similar sequences with increased developability through nonlinear dimensionality reduction and we explore the inferred developability landscape via nested sampling. The analysis enables the first direct visualization of the fitness landscape and highlights the existence of evolutionary bottlenecks in sequence space giving rise to competing subpopulations of sequences with different developability. The work advances applied protein engineering efforts by predicting and interpreting protein scaffold developability from a limited dataset. Furthermore, our statistical mechanical treatment of the problem advances foundational efforts to characterize the structure of the protein fitness landscape and the amino acid characteristics that influence protein developability.

**Significance statement:** Protein developability prediction and understanding constitutes a critical limiting step in biologic discovery and engineering due to limited experimental throughput. We demonstrate the ability of a machine learning model to learn sequence-developability relationships first through the use of high-throughput assay data, followed by the transfer of the learned developability representation to predict the true metric of interest, recombinant yield in bacterial production. Model performance is 44% better than a model not pre-trained using the high-throughput assays. Analysis of model behavior reveals the importance of cysteine, charge, and hydrophobicity to developability, as well as of an evolutionary bottleneck that greatly limited sequence diversity above 1.3 mg/L yield. Experimental characterization of model predicted candidates confirms the benefit of this transfer learning and in-silico evolution approach.

## Introduction

Engineered proteins have broad utility as therapeutics^1^, diagnostics^2^, targeted drug-delivery vehicles^3^, and as commercial products including industrial enzymes^4^ and agricultural processing catalysts^5,6^. Beyond the primary function (such as binding affinity or enzymatic activity), the utility of the protein is also dependent on the ability to be manufactured, transported, and stored while maintaining functionality. Commonly termed developability^7,8^, this often-overlooked property – quantified by stability, solubility, and production yield – is not typically assessed until late in the commercialization pipeline^9,10^. Late-stage developability failures: i) require substantial time for engineering or discovery of a new lead, ii) add avoidable costs which are often passed on to the consumer, and iii) prevent the immediate use of proteins that would otherwise benefit society^11^. The ability to predict protein developability and design beneficial mutations would ease the manufacturing process by reducing the experimental effort in selecting lead candidates for further evaluation^11,12^.

Predicting protein developability from amino acid sequence is nontrivial due to a myriad of factors: i) the combinatorial space resulting from twenty canonical amino acids possible at each position produces an exceedingly large sequence domain, ii) the sequence-developability landscape is believed to be rugged where a single mutation has the ability to eliminate functionality^13^, and iii) traditional developability assays often drastically under-sample the landscape due to experimental constraints^14^. The combination of these factors suggests the creation of a sequence-developability model, and the accurate determination of the most beneficial mutations will require advanced models and sampling techniques.

Recent advances in protein modeling suggest that machine learning possesses the ability to accurately predict functionality with sufficiently thorough and high-quality training data^15–17^. However, it remains unclear which embedding, or numeric representation, for proteins results in the most accurate and efficiently trained model. The traditional one-hot embedding for categorical variables creates a sparse embedding that lacks knowledge of physicochemical similarities between amino acids and is likely to result in poor performance^18,19^. Alternative approaches attempt to utilize precomputed amino acid properties, such as AAindex^20^ or structurally-based properties, such as non-polar surface area or contact density^21^, to embed sequences. However, determining the correct set of properties to use can lead to an exhaustive yet still incomplete search. An increasingly popular approach is to utilize an evolutionary-based model trained from homologs^22–24^. Nevertheless, properties that impact natural proteins (likely including primary function, natural mutational rates, and likelihood of experimental sampling) may not be the properties useful for assessing developability. As a result, we believe the most efficient method of training a sequence-developability model will use more direct experimental developability proxies, collected in high-throughput (HT), that can be transferred to predict traditional developability metrics.

In this study, we aimed to train and test a sequence-based model to predict the developability for variants of the protein ligand scaffold Gp2. While specific variants of this 45-49 amino acid protein scaffold have been shown to possess novel binding activity^25,26^, serve as a diagnostic in PET imaging^27^, and inhibit growth of breast cancer cells^28^, many variants still possess poor developability. In a prior study, a series of three HT assays – on-yeast protease resistance, expression as a fusion with a fragment of split green fluorescent protein (GFP), and modular insertion in split β-lactamase – were validated by mutual information and prediction of Gp2 variant yield (mg/L) via bacterial expression in two *E. coli* strains – T7 Express lysY/I^q^ (I^q^) and SHuffle T7 Express lysY (SH)^18^. Herein, we assess the ability to first train a sequence-based machine learning model to predict HT assay performance and transfer the **Dev**elopability **Rep**resentation (DevRep) to improve the accuracy in prediction of a traditional developability metric (Figure 1). After building a predictive model, we, i) analyze the learned sequence representation to identify factors driving developability, ii) use enhanced sampling techniques to explore and portray the developability landscape, and to identify high-yield variants, and iii) validate the findings experimentally showing that in-silico directed evolution can significantly outperform random mutagenesis.

**Figure 1.**
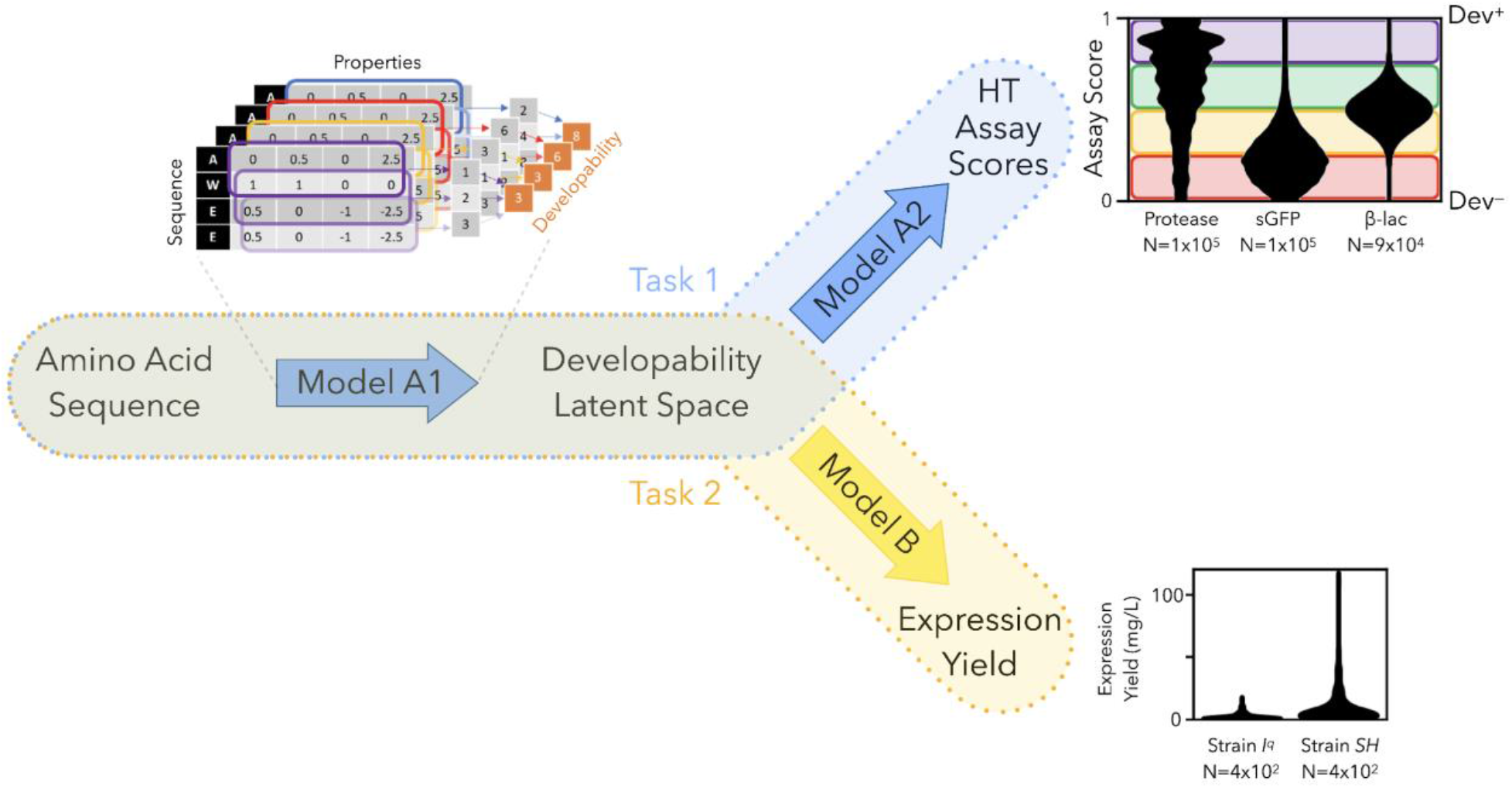
Prediction of protein developability via transfer learning. A sequence-based model to predict developability is trained in two steps. Task 1 (blue, top): The large database of protein assay scores is used to train a mapping (Model A1) from amino acid sequence to HT assay scores through a learned developability latent space representation (DevRep). Task 2 (orange, bottom): By transferring the representation, the expression yield (a traditional metric of developability) can be predicted when training a top model with a smaller dataset.

## Results

### Protein embeddings predict HT assay developability

A protein’s properties are determined by the interaction between amino acids, with various chemical properties, arranged in a three-dimensional structure uniquely determined by its linear amino acid sequence. We constructed models that first learn amino acid properties, and then combine them to create an embedding representative of Gp2 paratope variants (Figure 2a). We considered three architectures: i) *flat* - where all amino acid properties at all positions interact at once, ii) *recurrent* - where amino acid properties are fed one at a time into a memory unit that is updated as a function of the previously observed positions, and iii) *convolutional* - where amino acid properties are first summarized in a local region of the protein and then combined to obtain a full protein embedding. Multitask learning was applied to use all three HT assays to train a single developability embedding. Previous analysis revealed that this set of three HT assays was most informative and least redundant with respect to inferring recombinant yield, a low-throughput traditional metric for developability.^18^ We allowed dense layers between the protein embedding and assay scores (one to five layers permitted, hyperparameter optimization during cross-validation resulted in four layers) after the concatenation of a one-hot encoded assay-identifying vector.

**Figure 2.**
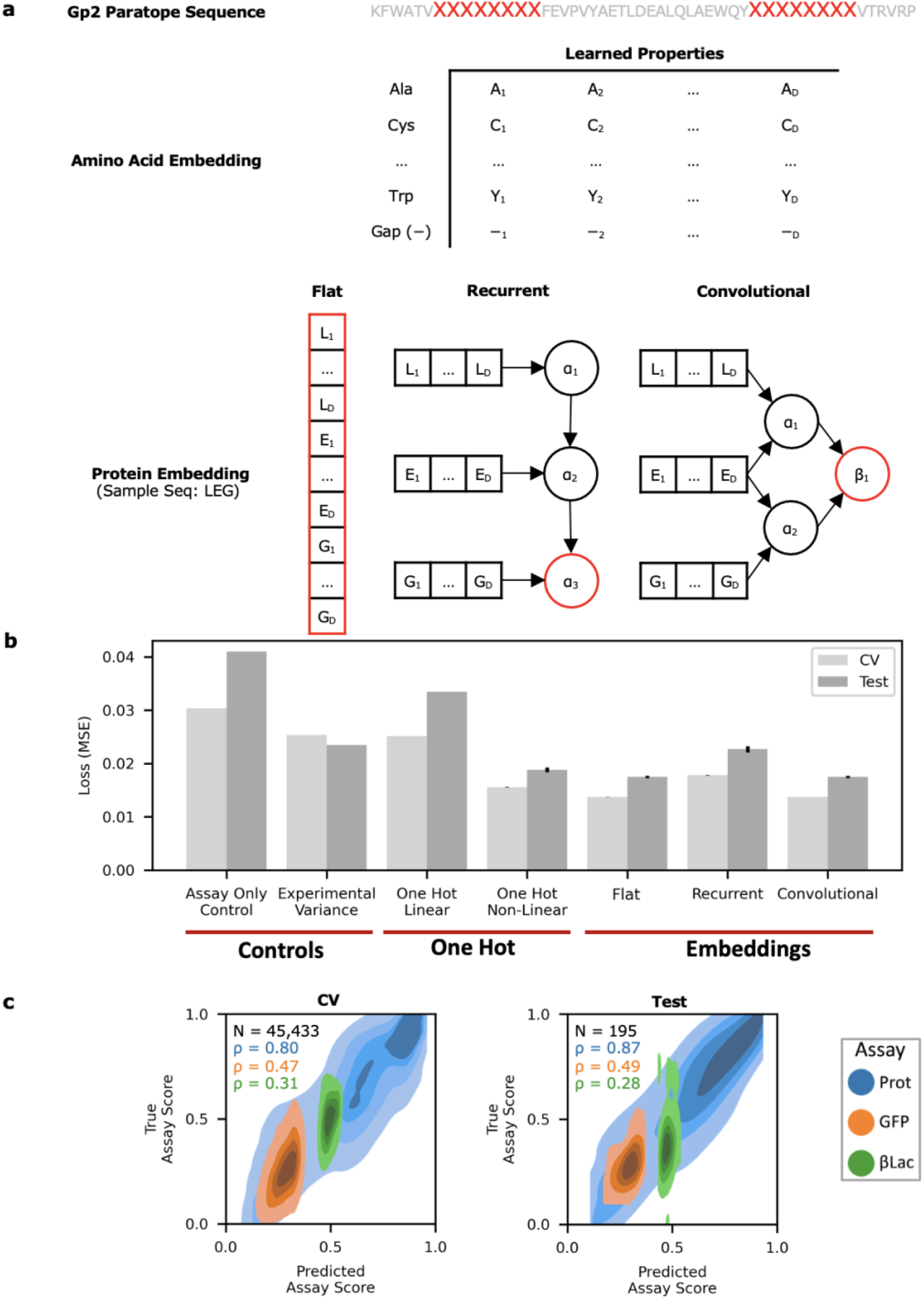
Protein embedding strategies based on interacting amino acid properties predict HT developability assay scores. **a)** The Gp2 paratope residues are embedded as learned amino acid properties and are combined via three different strategies into a developability representation, identified via a red outline. **b)** Embedded and non-embedded (one-hot) architectures were trained to predict assay scores via cross-validation (CV) and evaluated on an independent test set of sequences (independent 2-way Student’s t-test for embeddings vs. non-embeddings p<0.05). **c)** The convolutional architecture’s predictions are compared to the true assay scores (Prot: protease resistance, GFP: soluble expression in split-GFP system, βLac: modularity in split-β-lactamase) as a kernel density plot. The number of unique Gp2 variants and the Spearman’s rank correlation are displayed.

The performance of HT assay score prediction was compared to a series of controls as assessed by the mean squared error (MSE) of the cross-validation set and an independent test set (Figure 2b). All three architectures using sequence information were more accurate than the assay-only model (independent 2-way Student’s t-test p<0.0001 for all three embedding methods). We compared these architectures’ performance to the experimental MSE, *viz*. the variance of our measurements. This experimental assay variance was calculated as the sequence-averaged trial-to-trial (N=3) variance of the assay scores^18^. Dividing the variance by N=3 yields the squared standard error (SSE) (experimental assay CV SSE: 8.5 ×10^-3^; experimental assay test SSE: 7.8×10^-3^). The SSE represents our confidence in the assay scores when averaged over multiple observations/trials. Interestingly, the protein-inspired architectures can learn from multiple trials and thus predicts assay scores with lower MSE than the experimental variance (though not as low as the experimental SSE), highlighting the previously noted low resolution of a single trial of the assays^18^. We also compared the results to models that take as input a flat one-hot encoding of the amino acids of the Gp2 paratope (*i.e*., without linearly embedding the individual OH vectors first). We observe that a nonlinear model (flat sequence with dense layers between sequence and assay score) significantly outperforms a linear (ridge regression) model (independent 2-way Student’s t-test p<10^-6^). The FNN and CNN embedding models (which take in linear-embedding amino acid sequences as inputs) in turn significantly outperform the nonlinear model (independent 2-way Student’s t-test p<0.0001). We then visualized the relative correlation of the convolutional model’s predicted versus experimental assay score (Figure 2c). We found that the model was not equally predictive across assays, with the most accurate performance for the on-yeast protease assay.

### Testing Transferability to Traditional Developability Metric

Having developed a series of protein embeddings trained on and capable of predicting HT developability assay performance, we asked next if the same embeddings could be transferred to predict a traditional metric of developability. Keeping the embedding parameters constant (Figure 1, Model A1), we fit a separate top-model (Figure 1, Model B) to predict the soluble Gp2 yield in two *E. coli* bacterial strains – I^q^ and SH – via multitask learning using a one-hot encoded strain identifying vector. We used both linear (ridge regression) and nonlinear models (feedforward neural network (FNN), support vector machine (SVM) and random forest) to account for possible complex interactions between the embeddings and yield.

We found that transferring embeddings trained via assay scores resulted in the prediction of yield more accurately than a model trained directly from OH-encoded sequence to yield. During cross validation, the recurrent embedding with a random forest top model and the convolutional model with an SVM top model exhibited optimal performance (Figure 3a). Upon evaluation of an independent test set, the convolutional embedding with an SVM top model produced the most generalizable model (Figure 3b) while the recurrent embedding suffered from overfitting. Compared to the one-hot model with a random forest top model, the convolutional embedding reduced the gap to experimental variance (or MSE) by 44%. A Yeo-Johnson transformation was additionally individually applied to these yield measurements to remove correlation between error and yield^18^. The corresponding yield SSE divide the yield experimental variance by N=3 (experimental yield CV SSE: 0.117; experimental yield test SSE: 0.121). Additionally, the convolutional embedding was also able to outperform a model trained on experimentally measured assay scores (CNN test MSE: 0.53+/-0.03; experimental assay test MSE: 0.56 ± 0.004) (p < 0.05, independent 2-way Student’s t-test) suggesting the embedding can capture experimental assay information at least as well as a more traditional representation of the (experimentally determined) proxy HT assays to yield.

**Figure 3.**
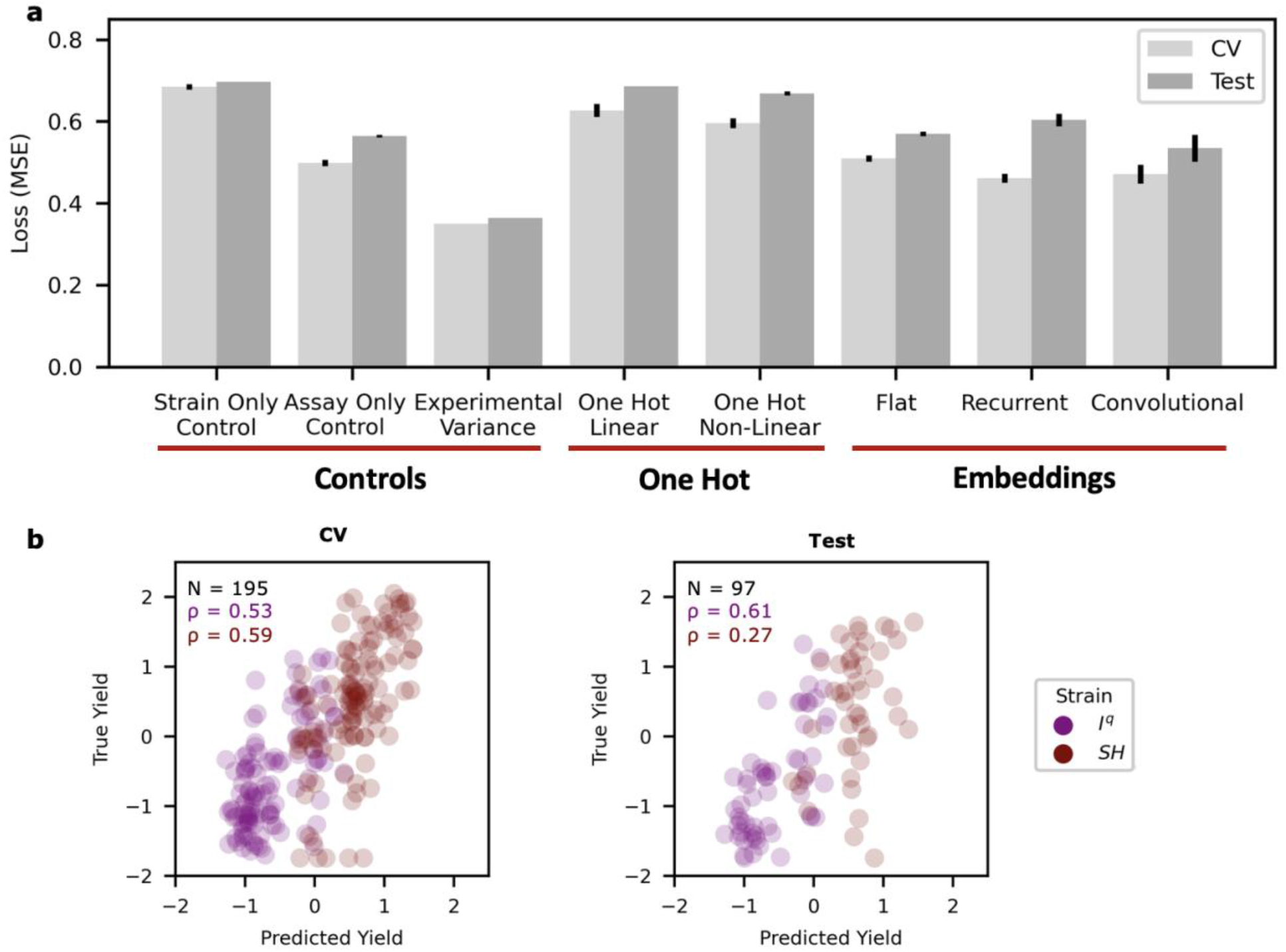
Transferred convolutional embedding predicts yield more accurately than traditional embedding strategy. **a)** Cross validation and Test performances of predicting yield comparing a traditional one-hot embedding to protein inspired embeddings trained by HT assay scores. **b)** The convolutional embedding with a support vector machine top model’s prediction of yields versus experimentally measured yield across *E. coli* strains I^q^ and SH.

### Dependence on HT Assays

Having observed the success of the transfer learning approach utilizing all three HT assays^18^, we sought to i) understand the importance of each individual assay in creating a transferable embedding and ii) understand if the transfer learning approach routinely creates a more informative representation than the direct use of HT assay scores. Each combination of HT assays was used to fit the three embedding architectures utilized in this study (flat, recurrent, and convolutional). The three top model architectures (ridge, random forest, SVM) were first trained on each HT assay combination’s embedding. The yield prediction accuracies of these models were then contrasted against those of the optimal top model (Figure 4a). The combination of all three HT assays created the optimal model. Combinations utilizing the on-yeast protease assay resulted in losses lower than those without (p < 0.01, independent 2-way Student’s t-test). In fact, this assay alone achieves an error within 2% of the model utilizing all three HT assays. This suggests that the on-yeast protease assay is the most informative assay and could potentially be used independently in future studies.

**Figure 4.**
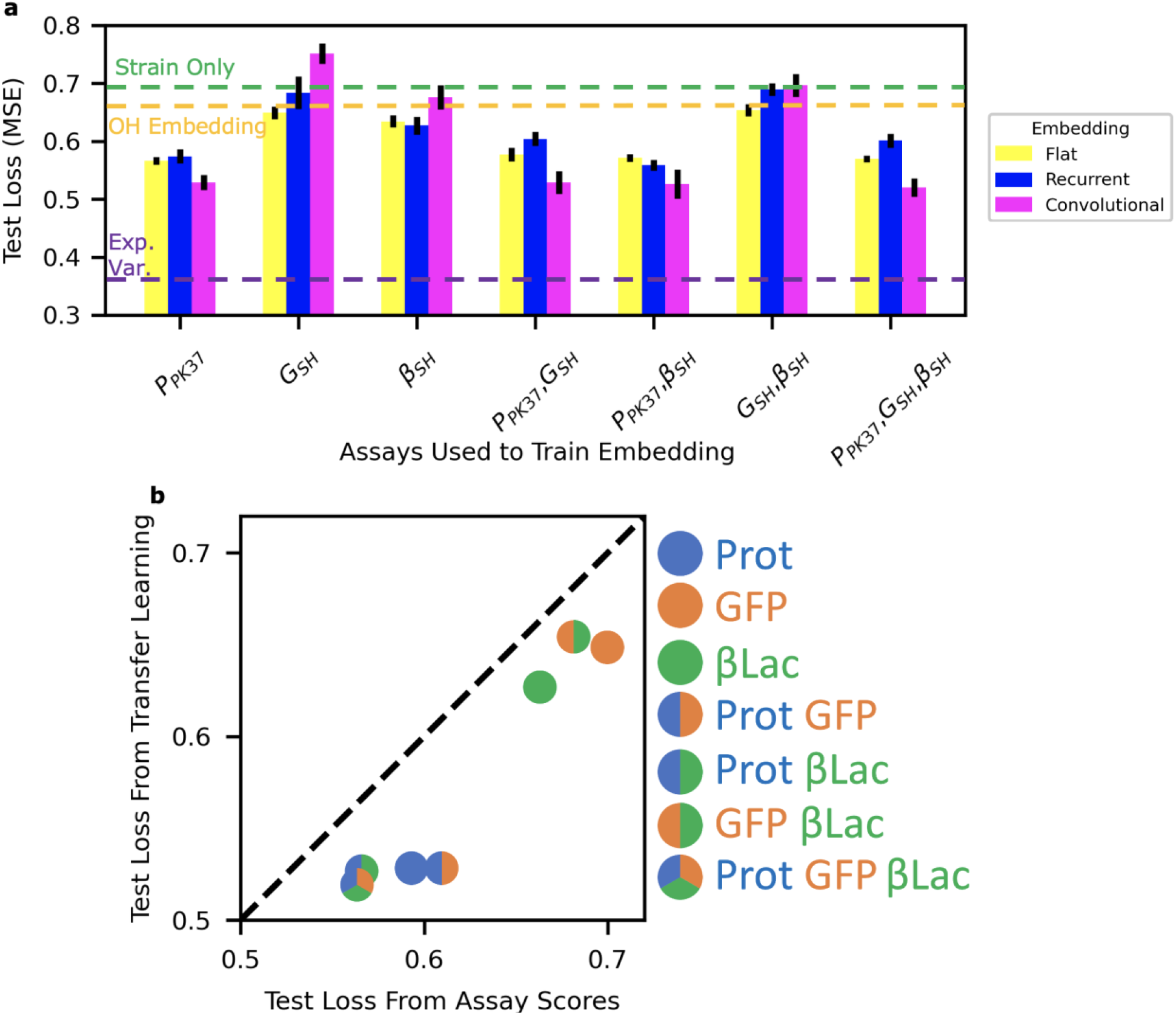
On-yeast protease assay is most informative and transfer learning enables discovery of true signal from imperfect HT assay proxies. **a)** A developability representation and top model to yield was trained with combinations of HT assays. The prediction error of sequence yield is grouped by assay combination and colored by embedding architecture. Error bars represent standard deviation of loss from N = 10 stochastically trained embeddings and top models. **b)** Yield predictions from assay scores and the most accurate trained embeddings for each combination of HT assays suggests transfer learning is more accurate than models that take as input the experimental assay scores.

The ability of the transfer learning training strategy to identify developability trends and average out noisy signals from similar sequences enables more accurate predictions than direct use of experimental HT assay outputs. Indeed, we previously observed the transfer learning model slightly outperformed a direct assay score to yield model (Figure 3a). We therefore sought to understand if transfer learning was successful because of the use of multiple assays and/or of the learning strategy more generally. To answer this question, we plotted the accuracy of models trained directly on experimentally measured assay scores to the accuracy of models that utilized the assay scores to train a representation that was transferred to predict yield (Figure 4b). We observed a correlation (Spearman’s ρ = 0.96) between the losses, suggesting that the more relevant assay score combinations enable more accurate embeddings regardless of learning strategy. We also observed a significant systematic decrease of loss from transfer learning models compared to models trained directly on assay scores (independent 2-way Student’s t-test p < 0.01), Fig. 4b. The ability of transfer learning to consistently outperform assay score models, even when a single assay is used, suggests the model can utilize sequence information to denoise errors present in the assay output.

### Alternative Model Building Approaches

After successful construction of the transfer learning model, we sought to validate the optimality of our transfer-learning strategy. In one alternative strategy, we evaluated the possibility of using DevRep to predict HT assay scores from sequences, and then to predict the experimental yield measurements from these predicted assays scores (rather than from the DevRep embedded sequences). In another, we evaluated the possibility of predicting yields from the experimental assay scores first, and then to train a sequence-to-yield model on these predicted yields (skipping the transfer learning altogether). We display the results of this analysis in Figure 5, showing DevRep controls (a), predicted high-throughput assays (b), and predicted yields (c).

**Figure 5.**
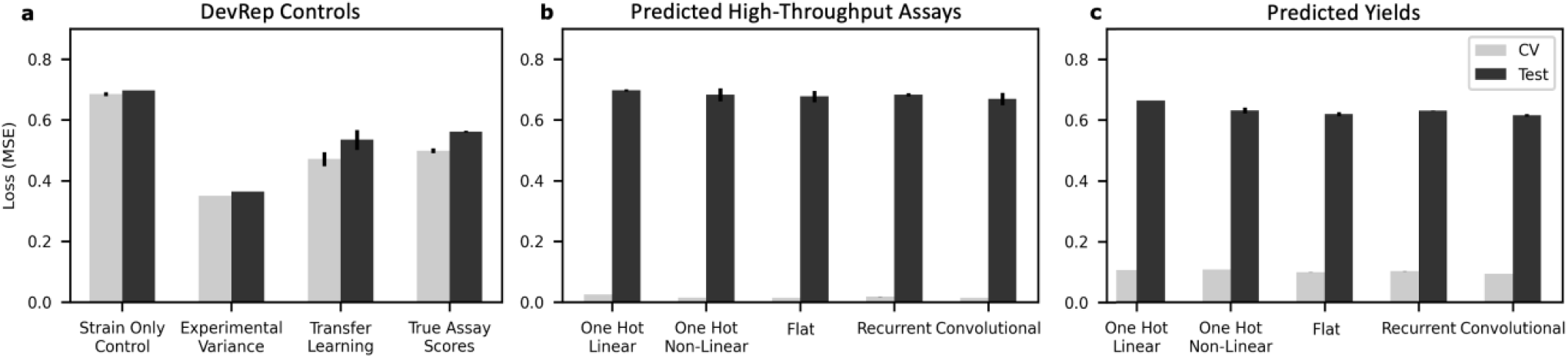
Alternative model cross-validation and test performance. **a)** DevRep controls (first outlined in Figure 3a). **b)** Predicted high-throughput assays are used to predict yield. **c)** Sequence-to-yield model trained on yields predicted from experimental HT data.

Briefly, both alternative strategies displayed significantly poorer performance and greater overfitting relative to the DevRep controls (Figure 5b,c). Further discussion of these regimes is available in Figure S1 and corresponding sections of Supplementary Text and Methods 1 and 1.4, respectively. This analysis supports our choice of developing a transfer learning approach in which only *experimental* (rather than predicted) assays and yields are used to learn a developability representation that when used as the input (*viz*., transferred) to a specialized model outperforms alternative approaches.

### Dependence on sample size

We next sought to understand the relationship between the size of the training datasets and the accuracy of the model. To this end, we randomly subsampled unique sequences from the HT assay dataset to develop convolutional embeddings and compare performance to non-transfer learning models that take as input a simple one-hot encoded representation of the sequence, or the experimental assay scores (Figure S2). DevRep performance systematically improved relative to controls as training data increased, highlighting further potential performance improvements from more data. The control models show greater efficiency on smaller sample sizes compared with DevRep.

### Model Interpretation

We have shown that an accurate model for the prediction of developability metrics such as the soluble yield of Gp2 in *E. coli* can be obtained by transfer learning. Specifically, we utilize a convolutional model to infer an embedding from a set of three HT assays, and feed (transfer) the embedding to an SVM (top) model to predict yield. We refer to this model as “DevRep”, and in what follows we explore the physical significance of the learned representation, and ascend the resulting developability landscape, yielding both a quantitative visualization of the landscape and a library of diverse and highly developable variants. The candidates of this library were then experimentally found to be produced in higher yield than sequences obtained by random mutagenesis.

### AA Embedding

First, we analyzed the trained amino acid embeddings to determine what properties are most relevant to the developability of Gp2 (Figure 6a). We evaluated if we could identify linearly separable latent amino acid features from our model via principal component analysis (PCA). The 17 feature dimensions (or inferred “properties”) were distilled down to three principal components (PCs) which explain 68% of the total variance. Upon inspection, we determined that cysteine is uniquely separated in PC 1 and 2. Additionally, PC 2 appears to separate the remaining residues by hydrophobicity by placing aromatic and aliphatic residues away from polar and charged residues. PC 3 further separates hydrophilic residues into negative, neutral, and positively charged. Interestingly, histidine (which possesses a pKa near experimental conditions) is located closer to neutral amino acids compared to arginine (R) and lysine (K), commenting on the ability of the model to learn about both charged states. We then compared each PC to the AAindex^20^ list of properties in an attempt to find the most correlative physicochemical property: PC 1 – coefficient over single-domain globular proteins (ρ = 0.91), a measurement of hydrophobicity^29^ again underscoring its importance on developability; PC 2 – normalized frequency of N-terminal non-beta region (ρ = 0.86), a measurement of residue frequency in nonstructured regions^30^; and PC3 – helix termination parameter at position j-2, j-1, j (ρ = 0.83), a measurement of residue frequency in short helical structures^31^. Together, PC 2 and 3 suggest that the paratopes may be balancing between a flexible loop and a short helical conformation to provide stability.

**Figure 6.**
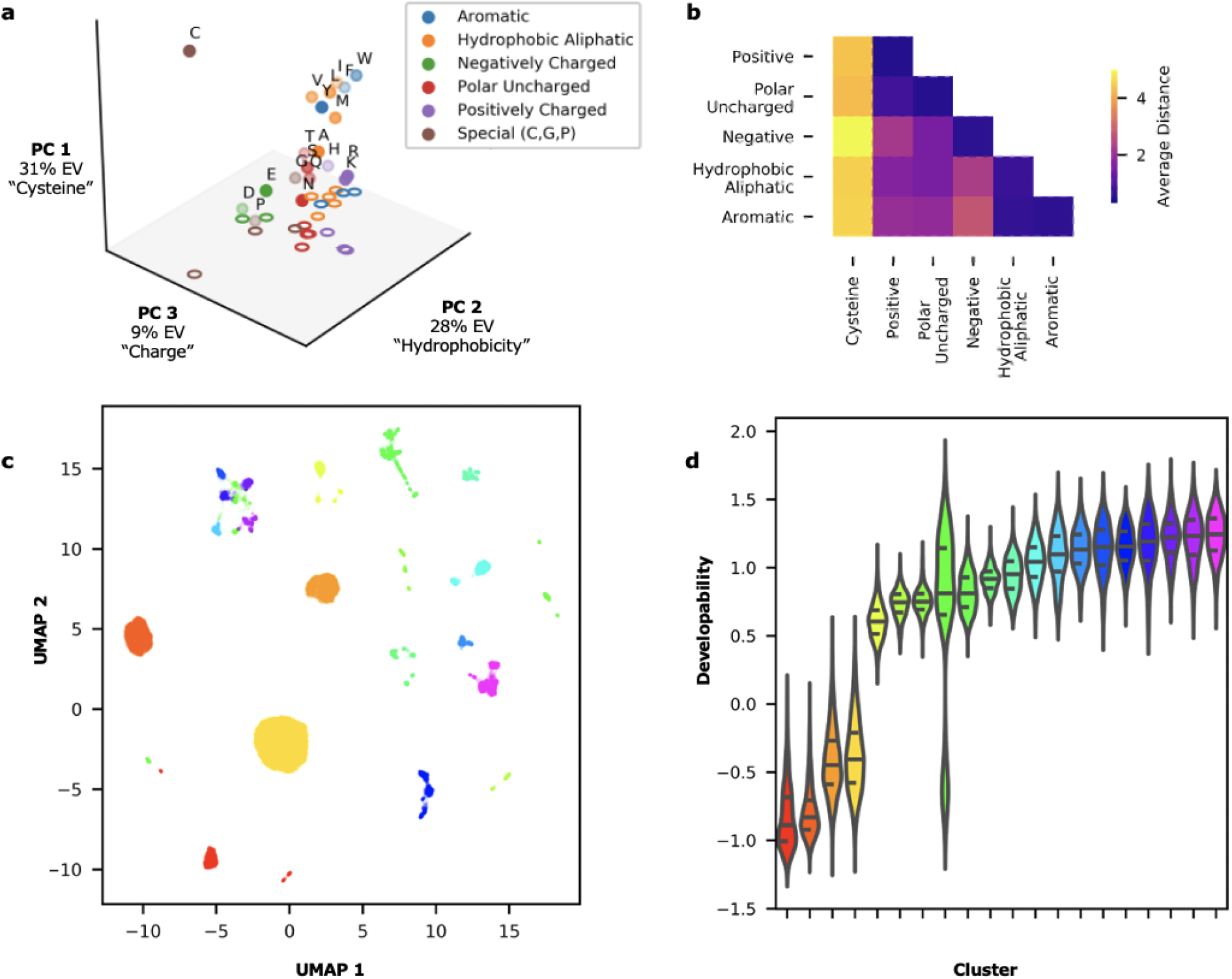
Analysis of trained embeddings reveals properties related to developability. **a)** Principal components (PC) of the 17-dimensional amino acid embedding, colored by category of residue. EV = explained variance. **b)** Inter- and intra-residue category distances highlighting the uniqueness of cysteine and lack of difference between aromatic and aliphatic residues. **c)** Clusters of sequences were identified via UMAP and hdbscan of the 45,000 sequences used for training. **d)** Developability, as predicted by yield, varies between clusters trained on HT assay scores.

We further evaluated the average inter- and intra-residue PC distances (Figure 6b). Each identified cluster of residues has a lower intra-residue distance than inter-residue distance except for aromatic (F, W, Y) and hydrophobic aliphatic (A, I, L, M, V) residues suggesting the hydrophobic nature of these residues outweighed the relative size difference and additional interaction capabilities of aromatic rings.

### Clustering of Training Sequences in Developability Space

We next assessed the interaction of the residue embeddings by converting the 97-dimensionial DevRep embedding for the 45,000 training sequences via UMAP^32^. UMAP accounts for nonlinearly related features; we thus chose UMAP (rather than PCA) for this analysis as we hypothesize that sequence embeddings are not linearly separable. We then utilized hdbscan^33^ to identify 19 clusters of sequences from the 2-dimensional UMAP space (Figure 6c). We discovered that the clusters contain information about the variant developability by finding a significant difference in developability distributions as a function of cluster (Figure 6d, Kruskal-Wallis H-test, p < 0.05). We further investigated the UMAP distributions of these clusters (Figure S3). This analysis suggests that poorly developable clusters (Figure 6d) are separated along a nonlinear manifold in DevRep space from highly developable clusters (Figure S3a). Additionally, the differences in amino acid frequencies between the selected clusters, paired with the amino acid embedding analysis, suggest that the model learned residues’ interactions, particularly with cysteines (Figure S3b).

### Comparison to Alternative Protein Embeddings

We next compared DevRep to models built with other state of the art protein embeddings. The AAindex^20^ was used to create an embedding based upon physiochemical properties. As the index is known to contain several similar entries, PCA was used to isolate 3 and 10 residue properties. The paratope was then converted into either of these sets of AAindex properties and flattened. We also compared DevRep to three embeddings trained on evolutionary properties: TAPE^23^’s transformer embedding which was trained on the Pfam^34^ database via predicting masked residues, UniRep^22^ which was trained autoregressively on the UniRef^35^ database, and evolutionarily (evo) tuned UniRep obtained by isolating homologous sequences to Gp2 via HMMER^36^ and updating the embedding via the Jax-UniRep^37^ software package. All evolutionary embeddings were tested by averaging over either the full sequence (“full”) or the paratope sequence (“paratope”).

Each model built on these embeddings was trained to predict yield utilizing the same architectures and hyperparameter search strategy as for DevRep (Figure 7a). We found that DevRep was able to predict yield significantly more accurately than every other embedding. Furthermore, the evolutionary based embeddings (particularly UniRep paratope) were able to predict yield more accurately than the strain-only and one-hot controls.

**Figure 7.**
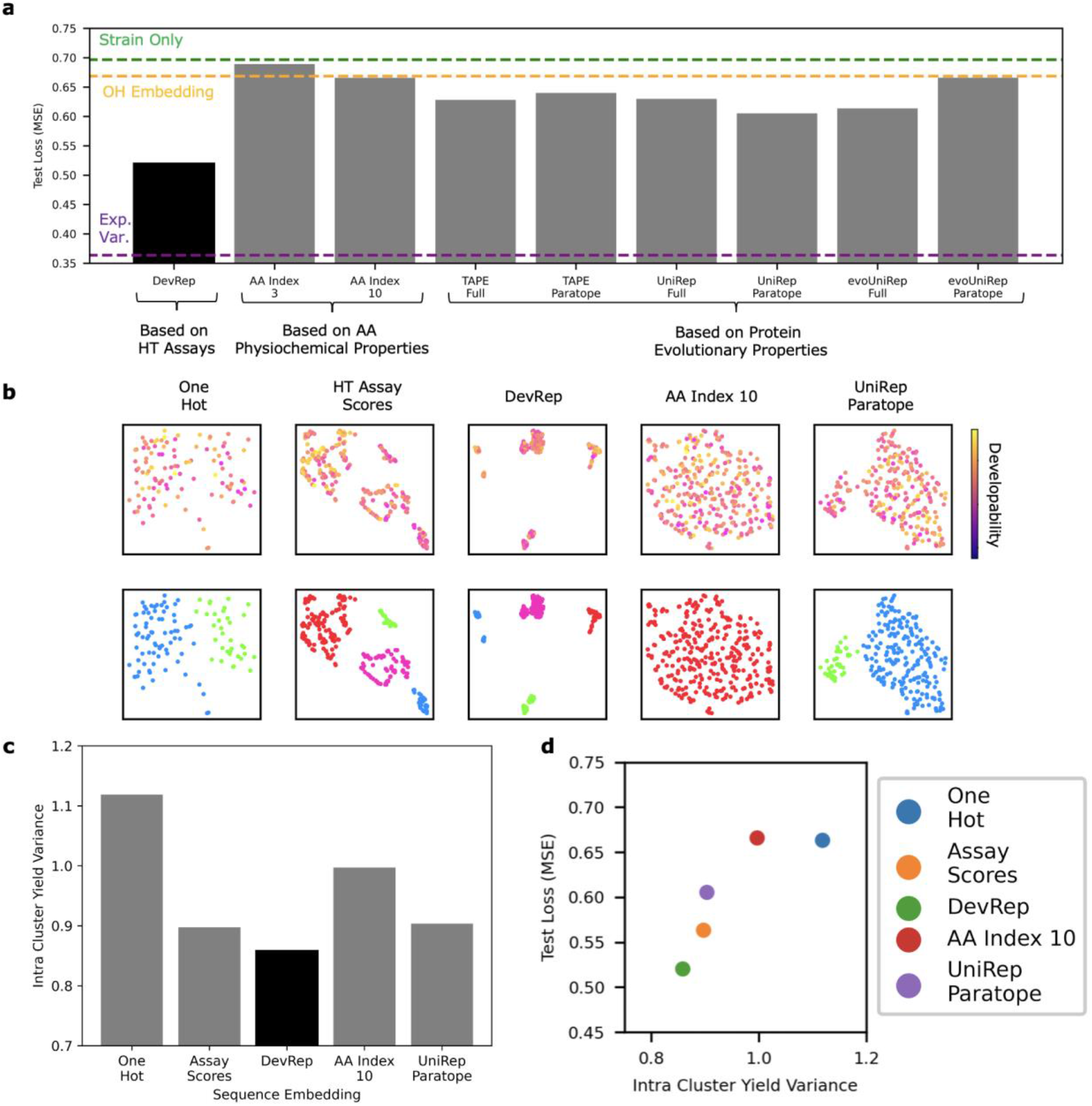
HT assay trained embedding contains more developability information than alternative embeddings. **a)** Comparison of protein embeddings’ ability to predict yield as represented by the loss on an independent set of sequences. **b)** Variants were plotted using UMAP for each embedding. (*top*) Color represents experimentally measured developability. (*bottom*) Sequences were clustered by UMAP coordinates. Color represents unique clusters. **c)** Variance in predicted yield across sequences within a given cluster. **d)** The correlation between the intracluster yield variance and the corresponding models’ (trained using the same embedding) predictive performance confirms that models that cluster sequences with similar yield also achieve better predictive performance, indicating that the embedding is informative about the predicted quantity (yield).

This realization highlights DevRep’s unique strength to find and use underlying developability relationships compared to other embedding controls. The poor performance of the AAindex also suggests traditional physicochemical properties are not the best way to describe Gp2 variant developability.

To ensure that the better performance of DevRep in predicting yield was not due to poor model development, we assessed the relationship between variation of sequences using each embedding and the measured developability. The 195 unique sequences with experimentally measured yield were embedded and transformed for visualization via UMAP (Figure 7b). We performed clustering in the UMAP space and calculated the average intra-cluster variance of the yield to estimate how much information about developability was encoded in the embedding; we associate lower transformed embedding intra-cluster variance with richer developability information content and presentation (Figure 7c). We found that the HT assay scores and DevRep’s UMAP representation was most informative about yield (Figure 7d).

### Sequence Space Analysis via Nested Sampling

Rather than rely on the skewed experimentally observed distribution of developability, we sought to use nested sampling to systematically characterize the structure of the fitness landscape while identifying highly developable sequences. At every iteration, nested sampling reduces the fraction of available sequence space “volume” (*viz*., the number of available sequences) by a constant proportion. Note that this sequence space is analogous to the more general phase space in statistical mechanics. As a result, we can use the output of nested sampling (a list of threshold sequences and their associated yield) to compute the density of states (DOS) as a function of developability (yield). Put simply, we can estimate the relative number of sequences available at any given developability (more generally any quantifiable fitness metric). Computation of the DOS also allows us to determine analogs of thermodynamic properties such as entropy, mean developability, and developability fluctuations (analogous to the heat capacity computed from the fluctuations in internal energy for a thermodynamic system). For example, in the context of developability, the analog heat capacity measures the rate of change in the mean fitness of the population upon varying selective pressure, ß. These thermodynamic analogs help to identify the occurrence of “phase transitions”. Such transitions occur when there are competing subpopulations of sequences with different developability, one of which becomes dominant upon varying selective pressure^38,39^. We ran the algorithm with 100 evolving sequences, removing the lowest yield sequence and thus contracting the phase volume by a factor of 100/101 (~0.99% of its original volume) at every iteration until convergence to a single sequence (Figure 8a). We then utilized the DOS to identify two main transitions: these transitions are apparent in the onset of relatively concave breakpoints corresponding to collapses in the DOS. Between these phase transition regimes, sequences split into multiple competing subpopulations at a given critical selective pressure β (an inverse temperature in a thermodynamic context). These transitions are highlighted by peaks in the heat capacity; the peaks correspond to high variance in sequence space as Gp2 variants transition from one basin of sequence space to another. The expected values of (β, developability) corresponding to these critical temperatures are (17.6, 1.3) and (46.5, 1.8), respectively (Figure 8b). This transition occurs with only 9.3×10^-5^ and 1.3×10^-10^ of all sequences predicted to have a higher yield, respectively.

**Figure 8.**
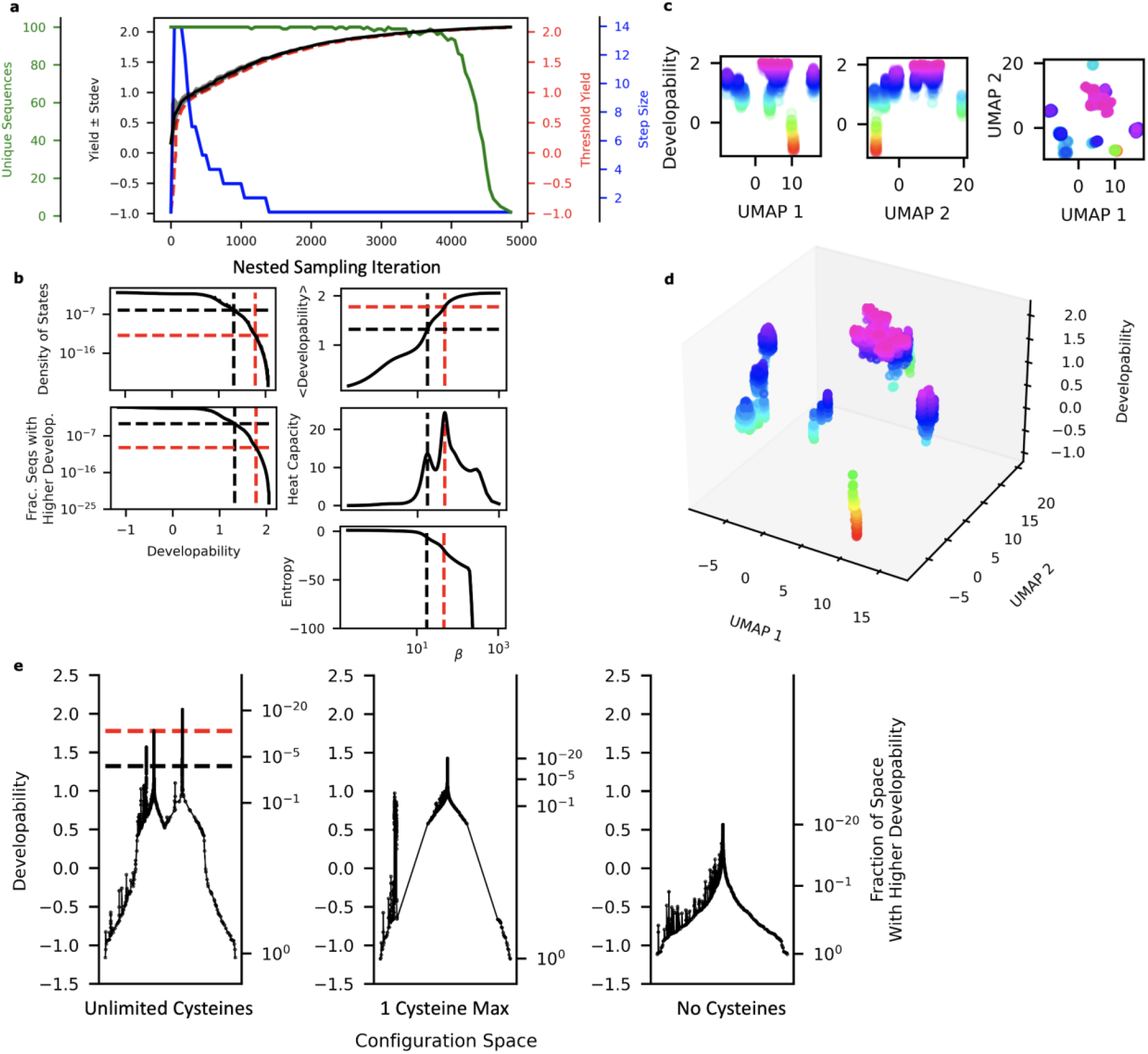
Nested sampling characterizes the developability-sequence landscape. **a)** Nested sampling was performed using 100 evolving sequences while accepting mutations with yields above the threshold per iteration. The threshold yield and corresponding sequences were determined by the lowest yield of the evolving sequences. **b)** The density of states for each level of developability was determined and used to estimate the expected developability, heat capacity, and entropy at various inverse temperatures (selective pressure in this context). Two main phase transitions are identified with a dashed line. **c,d)** The UMAP representation displays the landscape splitting into distinct clusters of DevRep space above the transition. Recorded sequences’ predicted developabilities increase from red to purple. **e)** The disconnectivity plot displays a landscape with competing developability peaks (when β grow large enough that a lower peak becomes depleted and a higher one enriched, we observe a phase transition).

The output of nested sampling can also be used to visualize the phase space^40,41^. Plotting the sequences in UMAP space shows a single stalk of low developability sequences up to the first phase transition where several high developability clusters exist (Figure 8c,d). The split suggests that beyond the first phase transition, there exists several distinct modes of achieving high developability. A disconnectivity plot was synthesized by creating a graph of nearest sequences of higher yield in the DevRep embedding^39–41^ (Figure 8e). The phase transition at the noted critical developabilities corresponds to a sharp decrease in available sequence space and branching occurs when subgraphs of sequences become disconnected at a critical developability level.

We compared disconnectivity plots and UMAP landscapes of the OneHot and UniRep Paratope models’ embeddings (Figure S4). Every model suggests at least one steep contraction of configuration space corresponding to a basin of similar sequences with high developability. The DevRep landscape is the only embedding to show a large split of sequence space into two competing basins. The OneHot UMAP landscape appears to have sequences of various predicted developability located at every UMAP location, confirming the one-hot embedding lacks easily interpretable developability information. The UniRep paratope landscape does show correlation between UMAP 1 and developability, suggesting there is some shared information between UniRep’s embedding space and developability space.

Previously we observed the importance of cysteine in distinguishing developability sequences according to DevRep’s embeddings (Figure 6). We hypothesized that restricting cysteine mutations within nested sampling would dramatically influence the resulting sequence-developability landscape. We thus further compared disconnectivity plots and UMAP landscapes of DevRep models’ embeddings when sampled sequences allowed for either i) at most one cysteine within a sequence or ii) no cysteine at all (Figure 8e). Indeed, we observe both the disappearance of the second basin within the disconnectivity plots and a significantly less developable final optimal sequence as cysteine is more stringently restricted.

### Identification of Top Developability Variants

As a final test of the transfer model approach to predict protein developability, we sought to measure the ability to predict high developability variants. Because we found that the Gp2 library splits into many subgroups of sequences that can achieve high developability (as it is also clearly visible from the multiple basins with high developability in the corresponding disconnectivity graph, see Figure 8e), we generated diverse sequences. We also identified sequences using simulated annealing^42^ to compare sampling strategies. The embeddings from each sampling approach were reduced via UMAP and clustered via hdbscan to identify sequences from clusters that are diverse in DevRep space. We then equally sampled across each cluster to acquire diverse high-yield variants. The most developable variants equally sampled in each cluster were recorded to yield a total of 100 final variants (Figure 9a, S5).

**Figure 9.**
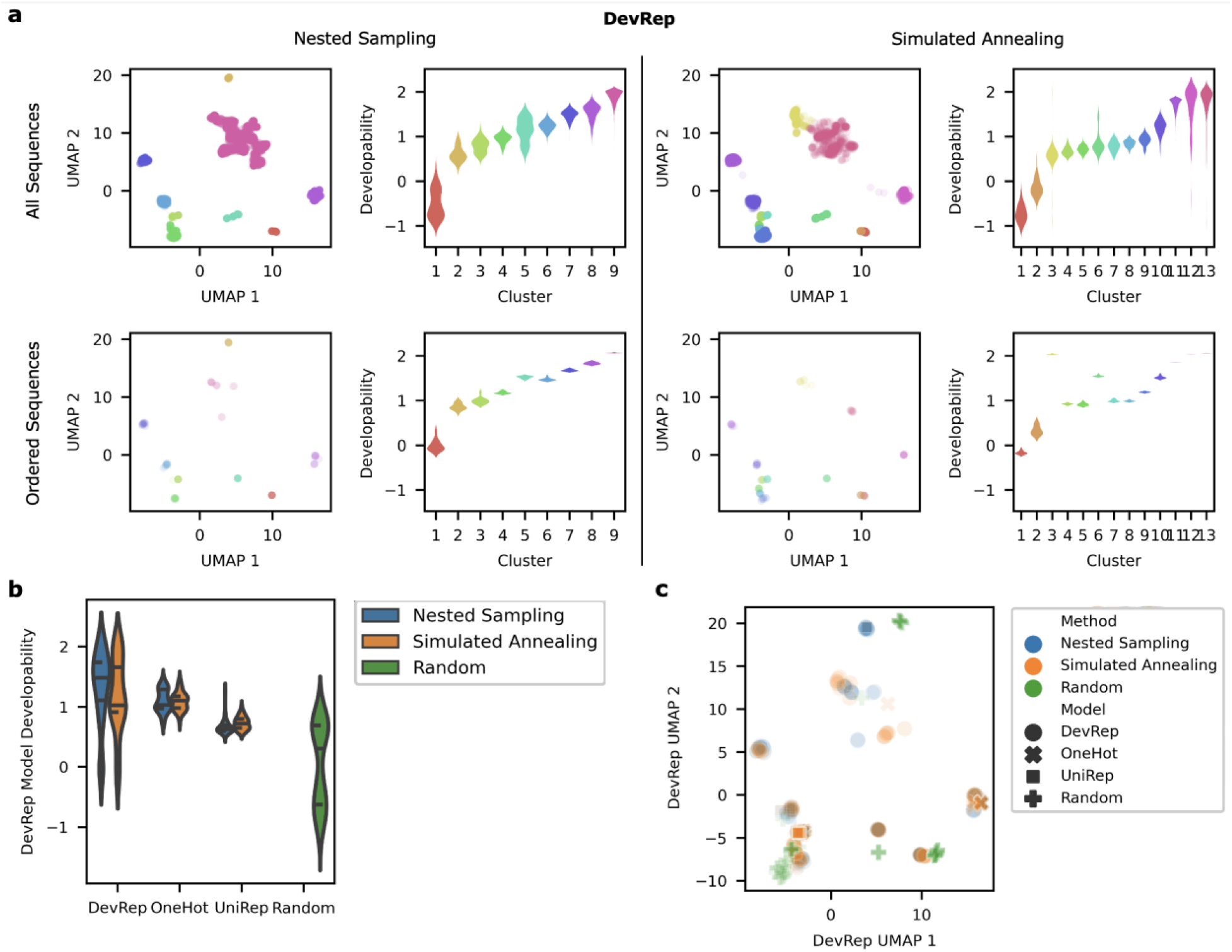
Assessment of DevRep-suggested high developability variants. **a)** Sequence embeddings identified through either nested sampling (left) or simulated annealing (right) strategies were clustered via UMAP (top) (Note: we only show the DevRep embedding here). The highest predicted yield variants in each cluster were equally sampled to determine 100 sequences. These variants represent a diverse set of sequences for experimental testing (bottom). **b)** Predicted developability distributions according to DevRep using equal inter-cluster sampling techniques across the sequence variants using different embeddings as in (a). **c)** UMAP visualization of top developability variants according to DevRep. Note that the UMAP visualizations of suggested top developability variants for nested sampling and simulated annealing in **a)** are shown in aggregate in **c)**.

The same process was repeated using different embeddings (OneHot and UniRep) for the paratope model. A randomly generated set of sequences was also tested for comparison. The predicted yields and different locations within sequence space for the isolated sequence embeddings suggest each model has its own distinct maximum and underlying approximation of developability space (Figures 9 and S5).

It was observed that including sequence diversity in the selection scheme introduced lower developability variants. Additionally, large clusters of high developability sequences were observed in both DevRep and UniRep embeddings from nested sampling (Figure S5). Thus, for each model, the large high developability cluster was split into subclusters where 100 additional variants were experimentally evaluated equally spread across the high-developability subclusters (Figure 10). 600 variants were thus proposed for experimental characterization.

**Figure 10.**
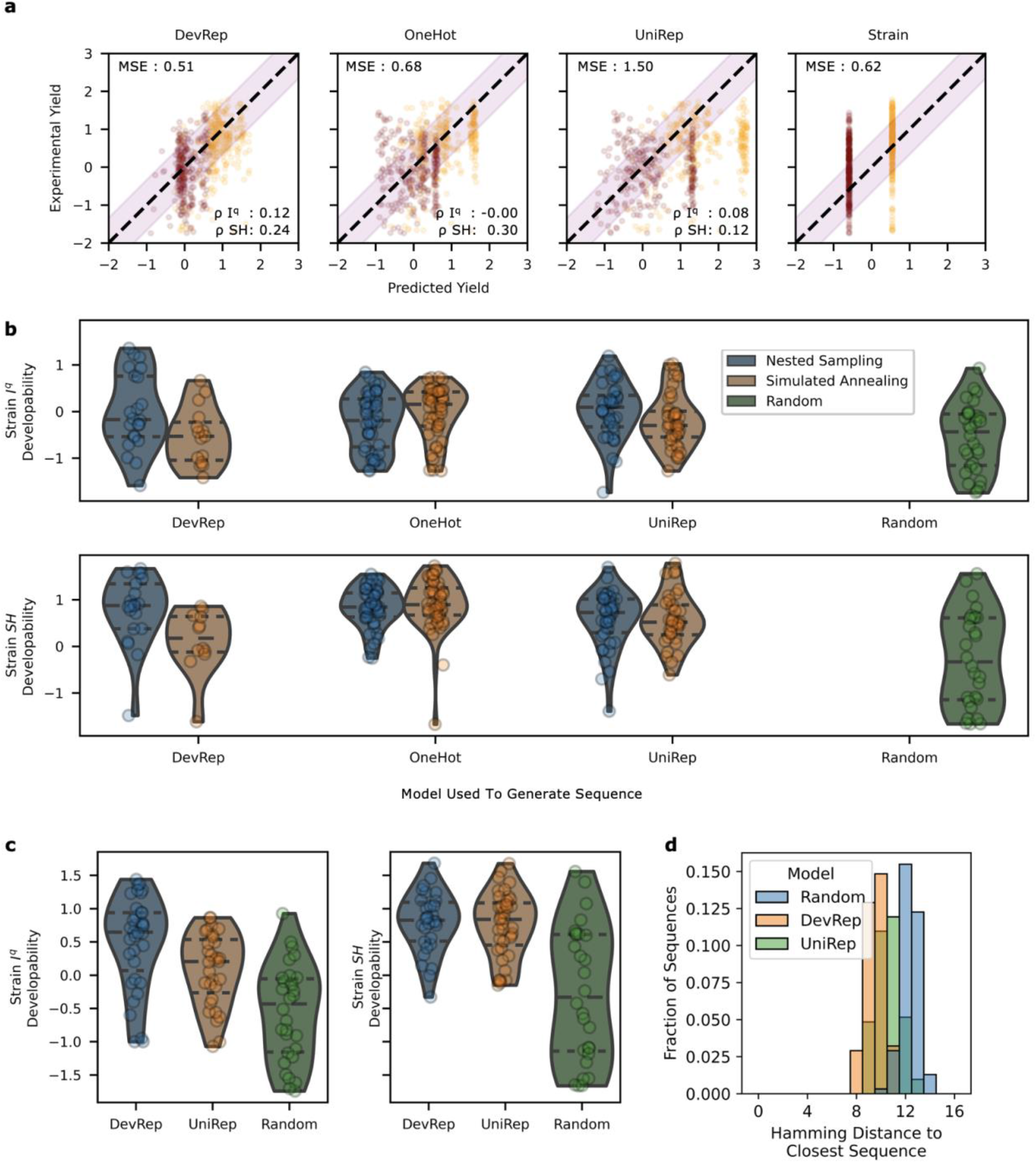
DevRep enables design of developable protein variants. **a)** The predicted versus actual developability of 280 I^q^ and 269 SH variants identified via sampling strategies (see Figures 4.12 and 4.13). **b)** Sequences generated by each embedding and sampling strategy are compared to each other and to a selection of randomly generated sequences. **c)** An additional set of sequences identified via nested sampling of DevRep and UniRep were also compared. These sequences were designed to be more developable and more similar in embedding space. **d)** Each sequence in (c) was compared to the set of sequences with measured yield that was used during model training. The distribution shown is broken down by the model used to generate the sequences.

We measured expression yield from the soluble fraction of *E. coli* bacteria for 280 variants in the I^q^ strain and 269 variants in the SH strain. The DevRep model was the most accurate in prediction of unseen sequences (Figure 10a). Interestingly, the OneHot encoded model outperformed the UniRep encoded model.

We next assessed which model and sampling technique identified the top performing variants with additional focus on diversity (Figure 10b). DevRep identified the highest yield sequences (upper quartile developability: 0.76 for DevRep vs. 0.27, 0.35, and −0.05 for OneHot, UniRep, and Random, respectively in strain I^q^ and 1.34 vs. 1.15, 1.01, and 0.61 in strain SH; Figure 10b). Nested sampling was more effective than simulated annealing for DevRep and UniRep but not OneHot embeddings in both strains.

We then assessed the distribution of yields obtained with a higher focus on developability than diversity (see Figure S6). Again, both DevRep and UniRep embeddings were able to select sequences with higher developability than a random selection (Figure 10c). Additionally, DevRep was able to identify the sequence with the highest developability in I^q^. Of final note, we found the sequences identified in this final evaluation were significantly far (in terms of Hamming distance) from variants evaluated during model training. DevRep’s sequences were 9.5 (on average) amino acid mutations away from the closest sequence during training (Figure 10d).

These results display a promising utility of DevRep in terms of both predictive accuracy and the ability to identify highly developable variants. Additionally, the performance of both OneHot and UniRep embeddings and sampling strategies suggest these techniques could be a useful first step in sequence identification, even prior to experimentation. Finally, we found that the combination of machine learning models with nested sampling constitutes a promising strategy for efficient and interpretable in-silico directed evolution of proteins.

## Conclusion

This work evaluated the ability of using HT developability assays (proxies for a traditional metric) to learn an embedding that is transferable to a predictive model of a traditional metric (*e.g*., yield) for which only few data points are available (an example of few-shot learning). We determined that this strategy can overcome noise in the proxy assays and achieve significantly better performance than alternatives. We then analyzed the model’s predictions to identify unique modes of achieving high developability based upon the location of cysteine and the importance of hydrophobicity and charge. The configuration space was explored via nested sampling which identified a range of developabilities where the sequences are highly clustered and unique, suggesting that a series of sub-libraries may outperform a single design. The transfer learning approach outperforms models based on physiochemical or evolutionary properties, thereby confirming developability is a complex and unique property and providing a combined experimental/bioinformatics means to integrate developability design into protein discovery and engineering.

## Acknowledgements

This work was support by the NIH (R01 EB023339 and R01 GM146372) and an NSF Graduate Research Fellowship (to A.W.G.). S. M. also acknowledges support from the Simons Foundation Faculty Fellowship and from the Simons Center for Computational Physical Chemistry at NYU, as well as partial support from the National Science Foundation (Award no. 2226387 and 2039575). We appreciate support from the University of Minnesota Flow Cytometry Core, University of Minnesota Genomics Center, and the Minnesota Supercomputing Institute at the University of Minnesota.

## Authorship and contributions

A.W.G., S.M. and B.J.H. designed the research. A.W.G., Z.D.S., B.J., S.M. performed the computational research. A.W.G., G.H.N. performed the experimental research. All authors analyzed the data. A.W.G., Z. D. S., G.H.N., B.J.H., S.M. wrote the paper.

