## Supplement for "Predicting and Interpreting Protein Developability via Transfer of Convolutional Sequence Representation"

**This PDF file includes:**

Supplementary text

Figures S1 to S6

Methods

### Supporting Information Text

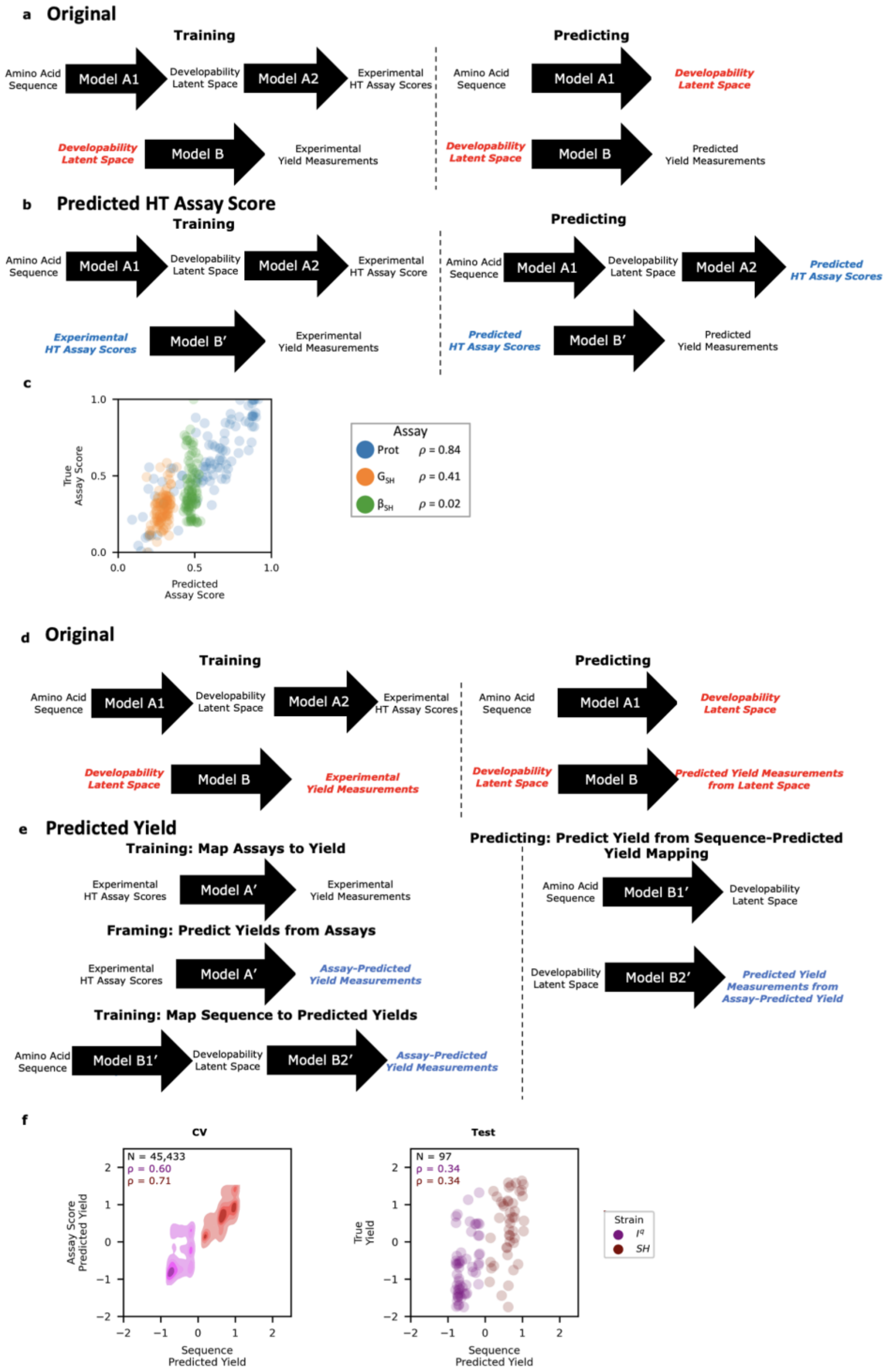


***Figure S1. Alternative model performance comparison.*** Two sets of architectures predicted either intermediate HT assay scores (a-c) or output recombinant yields (d-f). **a)** The final DevRep architecture consists of two sets of models that first construct a developability latent space mapping from amino acid sequence to experimental assay scores (Model A1 and A2) and then use this hidden state embedding to train a second model to predict yield (Model B). **b)** The predicted HT assay architecture instead predicts yield directly from predicted assay score (Model B'). **c)** Comparison of predicted versus actual assay scores for the convolutional embedding model. **d)** The final DevRep architecture makes use of a transfer learning step in which a hidden state serves as an input for a top model to predict recombinant yield. No predicted assay scores are used in the original yield prediction. **e)** The predicted output yield architecture instead generates a dataset of predicted yield measurements (Model A'). These predicted yield measurements are then used to predict yield from amino acid sequence (Model B1' and B2'). **f)** Comparison of the top-performing convolutional architecture performance on learning yields predicted from assay scores (left) and amino acid sequence predicted yields (right).

### 1. Alternative Model Building Approaches

First, we asked if predicted assay scores - generated using a sequence to assay score model - could be used to predict yield rather than using an intermediate latent space representation of the sequence trained on the experimental assay scores (Figure S1a-b). We found that the corresponding models failed to predict yield better than a strain-only control (Figure 5a). Upon further inspection, we observe that while the on-yeast protease assay scores were predicted well ($\rho = 0.84$), the split GFP ($\rho = 0.41$) and split β-lactamase assay ($\rho = 0.02$) scores were predicted incorrectly (Figure S1c). These predicted scores, compounded with the lack of sequence information, likely resulted in the poor yield prediction. The failure of this direct approach to predict yield from predicted assay scores suggests application of a sequence representation trained on experimental assay scores captures and frames developability information in a manner more suitable that using predicted assay scores.

We next asked if we could have fit a sequence-based model on the yields predicted from the experimentally measured assay scores instead of the smaller set of true yield measurements (Figure S1d-e). The set of 45,433 assay scores were converted to predicted yield in both bacterial strains and then used to train models with the same architectures as those employed for predicting assay scores (Figure 5c). All architectures displayed a strong ability to learn the yields predicted from the experimental assay scores, with cross validation losses ranging from 0.09 to 0.11. However, upon evaluation of the independent test-set sequences, all models displayed high levels of overfitting with test losses ranging from 0.62 to 0.66. We further observed that the top-performing convolutional architecture matched the yields predicted from assay scores (Figure S1f). This indicates that a sequence-based model trained on predicted yields across Gp2 variants is also insufficient to accurately generalize to predict the yield for Gp2 sequence variants.

### 2. Sample Size

The convolutional embeddings using 1%, 10%, and 100% of the HT assay data were transferred to the task of predicting yield (Figure S2a). Top models were then trained using 5%, 10%, 20%, 30%, 50%, and 100% of the yield training data (Figure S2b). At all fractions of yield data, performance was improved when training on more data suggesting that the larger HT assay dataset enabled the model to learn more universal developability information. Finally, we assessed the efficiency of our method by estimating how many unique sequences are required to achieve predictive accuracy within the experimental variance of yield measurements (Figure S2c). This experimental yield variance (measurement accuracy) was calculated as the sequence-average trial-to-trial (N=3) variance after applying a Yeo-Johnson transformation to individual trial yields[1]. Note that dividing this experimental variance by N=3 yields the average squared standard error (SSE) (Experimental Yield CV SSE: 0.117; Experimental Yield test SSE: 0.121). We found that the convolutional embedding trained by the full amount of HT assay data learns 90 ± 40 % more efficiently than the one-hot embedding only requiring $(0.6 \pm 1.7)\cdot 10^4$ unique sequences compared to the $(4 \pm 5)\cdot 10^4$ sequences required for the one-hot model. For smaller sample sizes, the higher MSE of the convolutional model compared to that of the OH model and the assay-only control (see Figure S2) indicate that the convolutional model tends to overfit at small training set sizes and compared to simpler embedding techniques like OneHot. In summary, more data and information always improve yield predictions, but basic, non-transfer learning embedding techniques are more economical in scenarios with data scarcity.

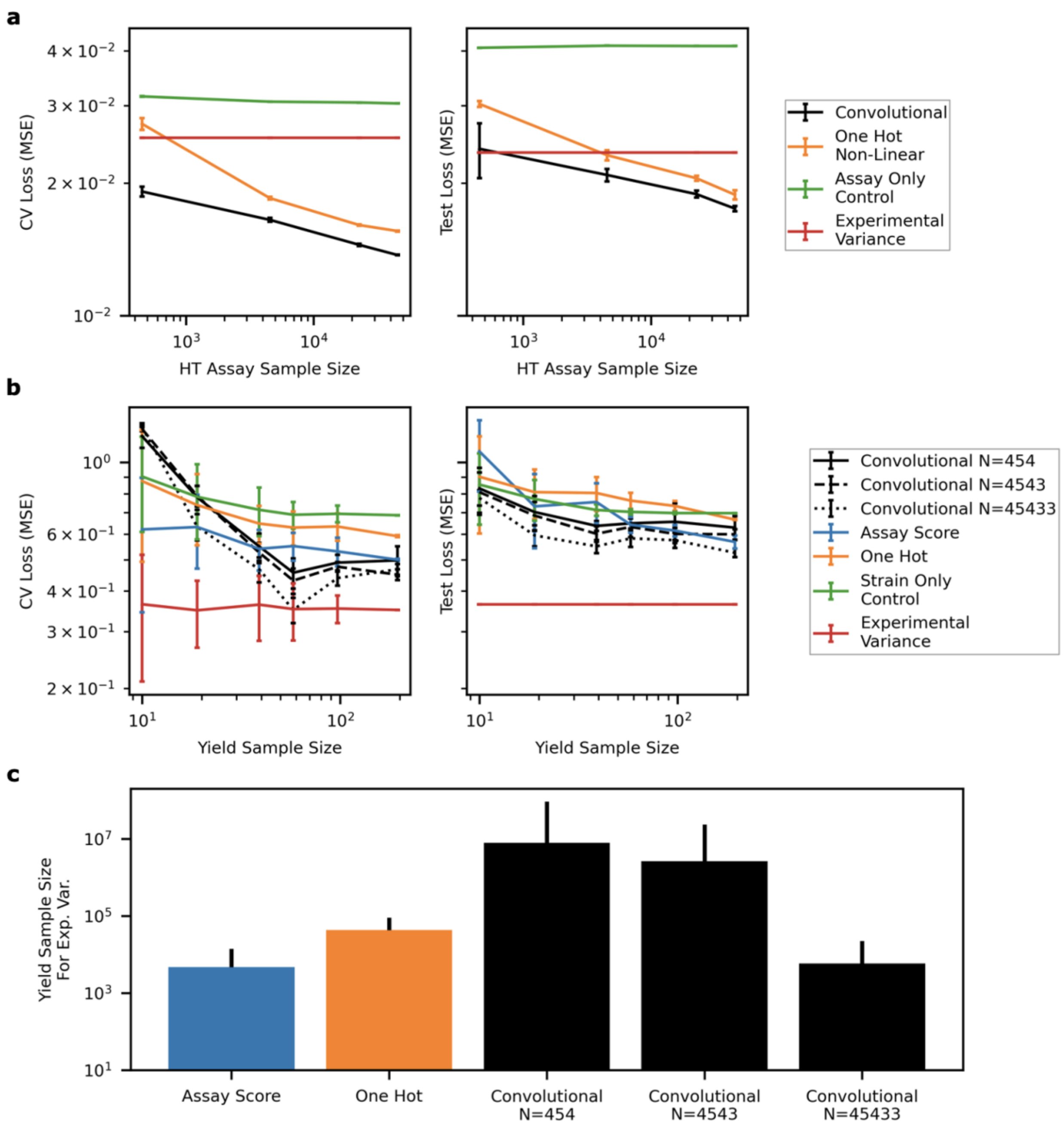


***Figure S2. Transfer model benefits from increase of sample size in both training steps.*** **a)** The convolutional embedding was trained on random subsets of the HT assay data. While performance improved with sample size, the relative performance over traditional embedding decreased. **b)** The convolutional embeddings from (a) were used to predict yield with top models trained via random subsets of available data. **c)** The predicted number of yield measurements to obtain a model with error matching experimental variance was extrapolated via log-log line of best fit weighted by the inverse of the confidence at each sample size. Error bars are propagated from the standard error of slope and intercept.

**3. AA Embedding**

Clustering and projection of our 45,000 DevRep-embedded Gp2 sequences via UMAP indicate the presence of distinct high and low developability clusters (Figure S3). For example, the location of the 4 lowest clusters were located close together in UMAP space (Figure S3a). Likewise, the higher-developability clusters all appear relatively further away from these lower-developability clusters. As UMAP can cluster sequences locally and globally[32], this suggests DevRep places most low-developability sequences in a similar location within the embedding. While we acknowledge UMAP provides no mathematical guarantees on inter-cluster distances being meaningful, we highlight that this global separation indicates that DevRep separates high- and low-developability sequences along a nonlinear manifold. We further analyzed the intra-cluster amino acid distribution for select clusters (Figure S3b): Orange - a low developability cluster not containing any cysteines, an increase in glycine, and loops of length 7; Yellow - a low developability cluster with cysteines at sites 7 and 12 with fewer glycines; Purple - a high developability cluster with cysteines at sites 7 and 12 and proline in the first loop; and Pink - a high developability cluster with cysteines at sites 7 and 12 and also in the second loop.

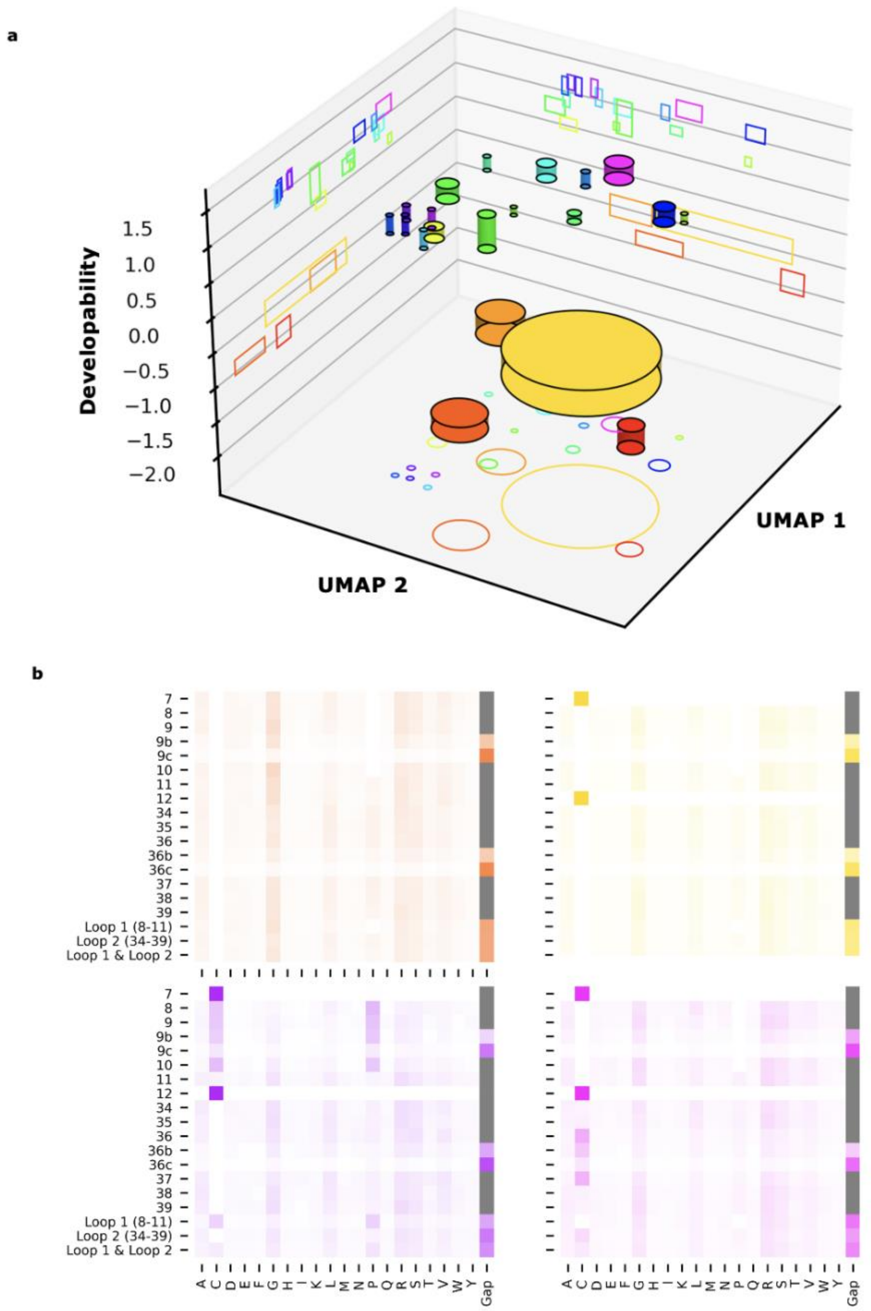


***Figure S3. Supplemental analysis of trained embeddings reveals properties related to developability.*** **a)** Three-dimensional landscape visualized by cylinders centered at the sequence-mean UMAP location with height representing the interquartile range of developability and radius corresponding to the number of sequences. Low-developability clusters are in the front right portion of the landscape. **b)** Amino-acid distribution of two low-developability clusters (orange and yellow) and two high-developability clusters (purple and pink).

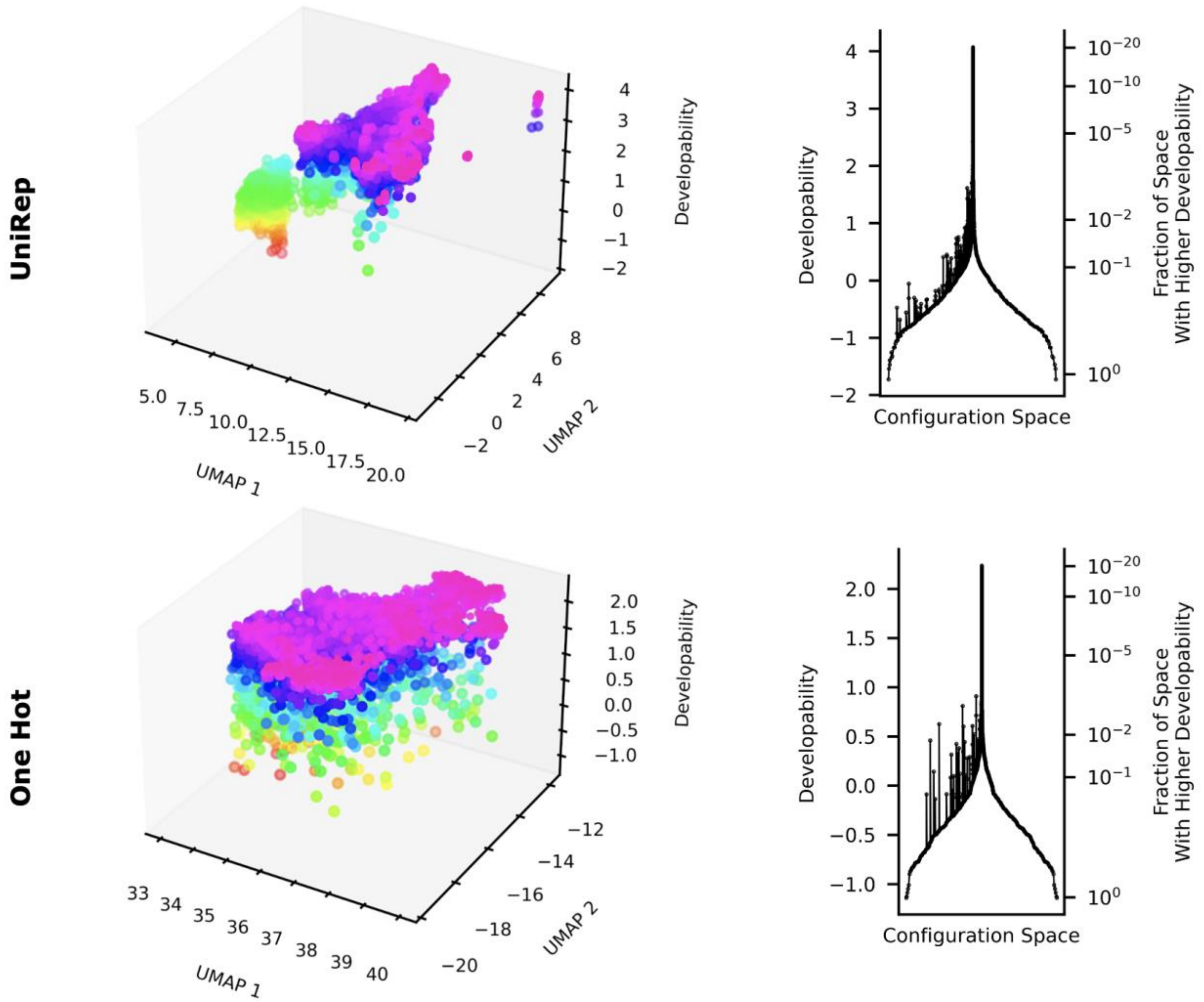


***Figure S4. Comparison of developability information contained across embeddings.*** Nested sampling was performed with each model. *(left)* UMAP representations of sequences sampled during sampling have varying relationships with predicted developability. Recorded sequences' predicted developabilities increase from red to purple. *(right)* Disconnectivity plots from different models agree with a sudden constraint in configuration space for the top 10% of sequences but disagree in the ability to identify a large split of sequence space.

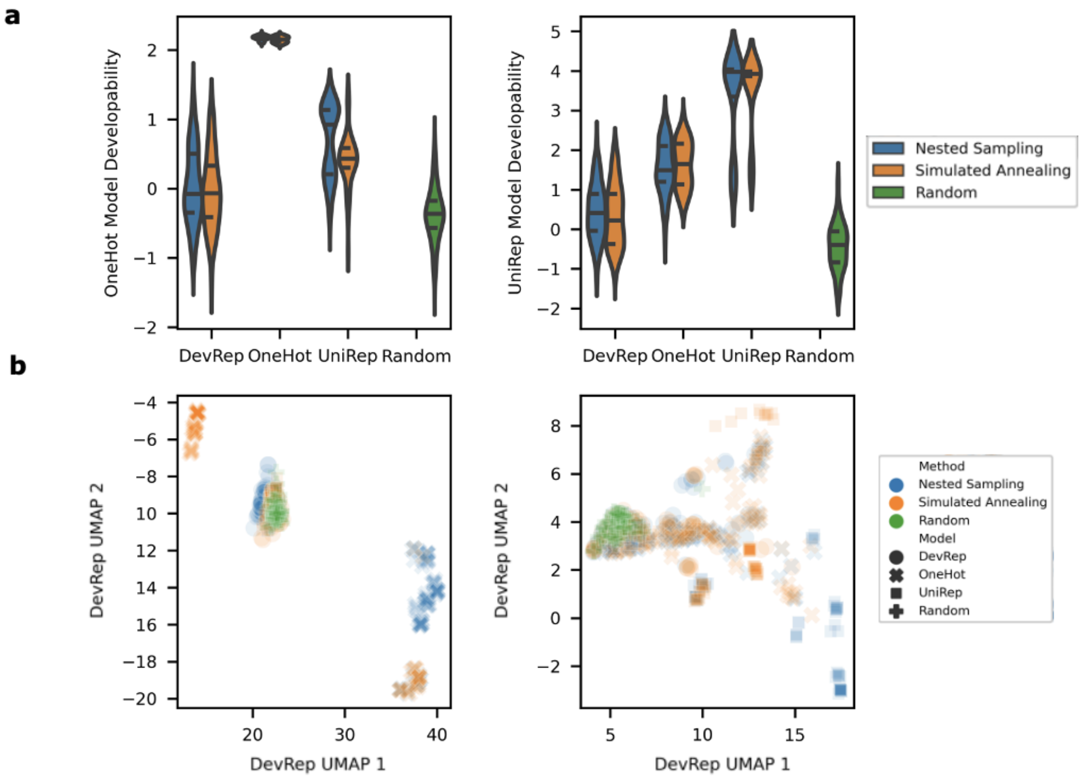


***Figure S5. Assessment of OneHot and UniRep-suggested high developability variants.*** **a)** Predicted developability distributions according to OneHot (left) and UniRep (right) architectures using sampling techniques across the sequence variants suggested by each architecture. **b)** UMAP visualization of top developability variants according to OneHot (left) and UniRep (right) architectures.

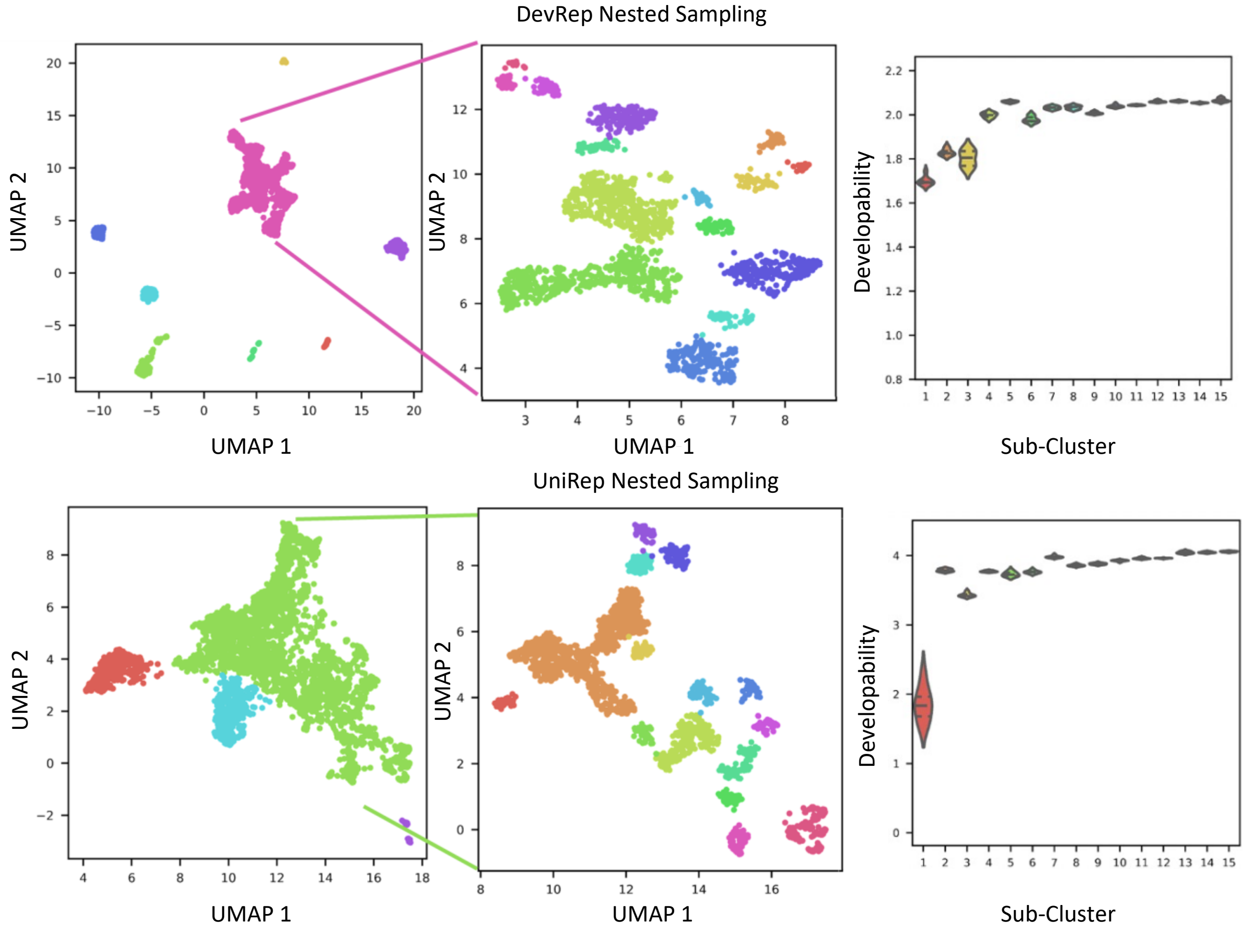


***Figure S6. Selection of additional high developability variants.*** Large clusters of highly developable variants were observed while clustering all recorded sequences during nested sampling. Thus, the clusters (left panel, DevRep - Purple, UniRep - Green) were broken into subclusters (middle panel) using the same methodology. The top sequences within each subcluster were selected for additional screening (right panel).

**Supporting Methods**

### 1. Model Development and Comparison

#### 1.1 *General*

All sets of examined architectures described here were developed in a similar format described below.

##### 1.1.1 *Code availability*

Python scripts used for deep sequencing and model evaluation, as well as datasets to train, evaluate, and plot performance are available at https://github.com/HackelLab-UMN/DevRep2.

##### 1.1.2 *CV performance*

Individual architectures were validated via either tensorflow or scikit-learn 10 times via k-fold cross validation (CV); embedding architectures were evaluated using tensorflow (k=3) whereas top models were evaluated using scikit-learn (k=10). The data within each assessed architecture set (*e.g.*, embedding strategy analysis) was conserved. Hyperopt determined the optimal hyperparameters for each architecture. Validation proceeded across either 50 trials or a maximum of 24 h of computational time; the trial with the lowest predictive error was recorded. The hyperparameters that resulted in optimal performance for each architecture within a set specified the best architecture for each architecture in a given set. Tables S1 and S2 summarizes the examined architectures across all sets with respective maximum assessed hyperparameter ranges.

##### 1.1.3 *Test performance*

All architectures within a set were retrained with their optimal hyperparameters on the entire CV training set. Held out test data for each architecture set was used to examine each architecture's performance. This independent test set was not used outside of examining architecture test performance.

**Table S1. Embedding architecture maximum assessed hyperparameter ranges**

| Architecture | Description | Maximum hyperparameter space |
|---|---|---|
| All | tensorflow.keras | epochs: (0,3), batch size: (0.1,1), number dense layers: (1, 5), dense nodes per layer: (1,100), weight dropout frequency: |

| | | |
|---|---|---|
| | | (0.1,0.5), amino acid embedding dimension size: (1,20) |
| OneHot | | |
| Flatten | tf.keras.layers.Flatten | |
| Recurrent | tf.keras.layers.Bidirectional, tf.keras.layers.GRU | input dropout frequency: (0.1,0.5) |
| Convolutional | tf.keras.layers.Conv1D | number filters: (1,100), kernel size: (1,16) |

**Table S2. Top-model maximum assessed hyperparameter ranges**

| Architecture | Description | Maximum hyperparameter space |
|---|---|---|
| Ridge | sklearn.linear_model.Ridge | alpha: (-5,5) |
| FNN | tensorflow.keras.layers.Dense | number dense layers: (1, 5), dense nodes per layer: (1,100) |
| Random Forest | sklearn.ensemble.RandomForestRegressor | number estimators: (1,500), max depth: 1,100), max features (fraction of total features): (0,1) |
| SVM | sklearn.svm.SVR | gamma: (-3,3), c: (-3,3) |

**Table S3. Sequence to Assay CV and Test Statistics (Corresponding to Figure 2)**

| Model | CV Loss Mean | CV Loss Error | Test Loss Mean | Test Loss Error |
|---|---|---|---|---|
| Assay Only Control | 3.03E-02 | 1.25E-07 | 4.10E-02 | 0.00E+00 |
| Experimental Variance | 2.53E-02 | 0.00E+00 | 2.35E-02 | 0.00E+00 |
| One Hot Linear | 2.51E-02 | 1.63E-06 | 3.35E-02 | 0.00E+00 |
| One Hot Non-Linear | 1.56E-02 | 3.92E-05 | 1.88E-02 | 4.55E-04 |
| Flat | 1.37E-02 | 1.29E-05 | 1.75E-02 | 2.24E-04 |
| Recurrent | 1.78E-02 | 4.47E-05 | 2.27E-02 | 5.86E-04 |
| Convolutional | 1.37E-02 | 2.49E-05 | 1.75E-02 | 2.33E-04 |

**Table S4. Sequence to Yield CV and Test Statistics (Corresponding to Figure 3 and Figure 5a)**

| Model | CV Loss Mean | CV Loss Error | Test Loss Mean | Test Loss Error |
|---|---|---|---|---|
| Strain Only Control | 6.85E-01 | 6.51E-03 | 6.97E-01 | 0.00E+00 |
| Assay Only Control | 4.99E-01 | 7.94E-03 | 5.65E-01 | 3.56E-03 |
| Experimental Variance | 3.50E-01 | 0.00E+00 | 3.64E-01 | 0.00E+00 |
| One Hot Linear | 6.27E-01 | 1.59E-02 | 6.85E-01 | 0.00E+00 |
| One Hot Non-Linear | 5.96E-01 | 1.17E-02 | 6.69E-01 | 5.04E-03 |
| Flat | 5.10E-01 | 8.01E-03 | 5.70E-01 | 5.76E-03 |
| Recurrent | 4.62E-01 | 1.10E-02 | 6.04E-01 | 1.55E-02 |
| Convolutional | 4.71E-01 | 2.28E-02 | 5.34E-01 | 3.30E-02 |

**Table S5. Predicted Assay to Yield CV and Test Statistics (Corresponding to Figure 5b)**

| Model | CV Loss Mean | CV Loss Error | Test Loss Mean | Test Loss Error |
|---|---|---|---|---|
| One Hot Linear | 2.51E-02 | 1.63E-06 | 6.97E-01 | 3.02E-03 |
| One Hot Non-Linear | 1.56E-02 | 3.92E-05 | 6.83E-01 | 2.25E-02 |
| Flat | 1.37E-02 | 1.29E-05 | 6.77E-01 | 2.10E-02 |
| Recurrent | 1.78E-02 | 4.47E-05 | 6.81E-01 | 4.03E-03 |
| Convolutional | 1.37E-02 | 2.49E-05 | 6.69E-01 | 1.94E-02 |

**Table S6. Sequence to Predicted Yield CV and Test Statistics (Corresponding to Figure 5c)**

| Model | CV Loss Mean | CV Loss Error | Test Loss Mean | Test Loss Error |
|---|---|---|---|---|
| One Hot Linear | 1.05E-01 | 5.56E-05 | 6.64E-01 | 0.00E+00 |
| One Hot Non-Linear | 1.07E-01 | 2.57E-04 | 6.32E-01 | 9.89E-03 |
| Flat | 9.86E-02 | 3.72E-04 | 6.19E-01 | 7.11E-03 |
| Recurrent | 1.02E-01 | 3.75E-04 | 6.30E-01 | 1.42E-03 |
| Convolutional | 9.32E-02 | 5.35E-05 | 6.16E-01 | 4.26E-03 |

**1.2** ***Protein embeddings predict HT assay developability***

All Gp2 sequences were either one-hot or ordinally encoded. Only the Gp2 paratope of length 12-16 amino acids was encoded; each paratope was padded with gap characters in positions 3,4 and 11,12 as needed to obtain a 21 (20 canonical amino acids plus a gap character) x 16 (positions) input matrix (for one-hot) or a 1x16 input vector (for ordinal). Combinations of embedding architectures (Table S1) and all top-models (Table S2) were trained, validated, and tested on appropriate splits of the 45,433 Gp2 variants following Section 1.1 to simultaneously predict the three optimal developability assays determined previously[1].

**1.3** ***Testing Transferability to Traditional Developability Metric***

The exact paratope sequence embeddings learned in Section 1.2 were then used as input to train all top-models (Table S2) to predict recombinant yield across the two examined Gp2 strains $I^q$ and SH. The cross validation (N=195) and test (N=97) partitions of the Gp2 recombinant yield dataset were used to assess these top models as described previously[1].

**1.4** ***Alternative Model Building Approaches***

The predicted assay score and predicted yield architectures were assessed similarly to Sections 1.1-1.3 with notable exceptions described below. The core motivation between these alternative regime investigations were to either 1) Predict intermediate information via training with predicted assay scores, or 2) Predict output information via training with predicted recombinant yields.

**1.4.1** ***Predicted HT Assay Score***

The predicted HT assay score regime overview is given in Figure S1a,b. All combinations of embedding models A1 and a subset of top-models A2 were trained, validated, and tested according to Section 1.1-2 to generate 45,433 Gp2 (sequence, *predicted high throughput assay*) datapoints for the three assays used previously[1]. The subset of top-models (Table S2) here was {Ridge (One Hot Linear), FNN (One Hot Non-Linear)} due to the other top-models' slower training and demonstrated poorer performance on this task relative to the tested top-models. This predicted high throughput assay dataset was then used as input to train a separate top model B' to predict recombinant yield using the N=195 and N=97 validation and testing datasets, respectively. This model B' was relegated to the Random Forest top model of Table S2 due to its previous success in identifying the optimal developability assays to predict recombinant yield[1].

**1.4.2** ***Predicted Yield***

The predicted recombinant yield regime overview is given in Figure S1d,e. Random Forest top model A' was trained, validated, and tested according to Section 1.1-2 to generate the 45,433 Gp2 (sequence, *predicted recombinant yield*) datapoints for the three assays used previously[1]. Model A' was assigned to the Random Forest due to its previous success in identifying the optimal developability assays to predict recombinant yield as in Section 1.4.1.

This predicted recombinant yield dataset was then used as input to train all embedding architectures in Table S1 (Model B1') with all top models in Table S2 (Model B2') to predict these

45,433 predicted (rather than the 195+97=292 experimental) recombinant yields per Section 1.1-1.2.

#### 1.5 *Dependence on sample size*

To explore the dependency of the top models' performance on embeddings trained over a range of sample sizes, all 45,433 Gp2 sequences were first converted into DevRep's convolutional embeddings. These 45,433 (embedding, assay score) datapoints were then subsampled at 1%, 10%, and 100% of the full dataset to train and validate all top models from Table S2 to predict recombinant yield. The held out (validation) and test performance on these dataset partitions was recorded. These performances were then compared to one-hot and assay-only embedding controls (Figure S2a).

Similarly, each of the top models in Table S2 was trained on 5%, 10%, 20%, 30%, 50%, and 100% of the N=195 datapoints for which all three assay scores and recombinant yield were available and their corresponding validation and test performances were recorded alongside appropriate embedding controls (Figure S2b).

Lastly, linear regression was performed on the number of training sequences against test loss for all combinations of assessed inputs {sequence, high throughput assays} and top models from Table S2. The intersection of this linear approximation with experimental variance was recorded (Figure S2c).

#### 1.6 *Dependence on HT Assays*

All combinations of embedding architectures (Table S1) and top models (Table S2) were trained on all combinations of the three assay scores to predict recombinant yield following Section 1.1-1.2 (Figure 4a).

A separate set of top models (Table S2) were then separately trained to predict recombinant yield from all combinations of the three assay scores, omitting any sequence information. The best top model's performance for every assay combination was recorded. Then, the best embedding architecture-top model combination across each assay combination (Figure 4a) was recorded to assess transfer learning's benefit to using the developability information inherent in each combination of assays. These two values of best performance for every assay combination were then compared (Figure 4b).

### 2. Model Interpretability

#### 2.1 *Amino Acids*

After DevRep validation and retraining, all 20 amino acids' learned features were concatenated to yield a 20x17 matrix. Principal component analysis on this matrix was performed via scikit learn to yield the three PCs that explained 68% of the total variance across amino acid representations. We compared these PCs against the physicochemical properties for each amino acid within the AAIndex library via pyaaisc: the PC-AAindex pair across all amino acids with the highest spearman’s correlation (calculated via scipy) was recorded. Inter- and intra- PC category were calculated via Euclidean distance using scipy cdist.

### 2.2 *Sequences*

2D UMAP-transformed sequences (via umap) had their components clustered via hbdscan with minimum cluster size of 1% of total sequences in the dataset; all nonclustered sequences were removed in later analysis. Each embedding (DevRep, OneHot, and UniRep) served as input to its distinct UMAP object with default parameters including n_neighbors=15, n_components=2, and min_dist=0.1 with a Euclidean distance metric. Note that no tuning of UMAP transformation parameters was attempted; our analysis instead focused on examining the global structure of projected embeddings and their associated developabilities. Resulting cluster representation distributions were compared with an H-test via kruskal from scipy.

### 2.3 *Sampling Procedures*

A set of 100 paratope sequences – known as walkers – of length 16 were initialized via random selection of all 20 amino acids at every position from a uniform distribution. Gap insertions were further allowed at positions 4-5 and 12-13 to enable encoding of paratopes of total lengths from 12 to 16 amino acids while yielding consistently 16-character sequences. If relevant, the assignment of cysteine to a position was restricted during this initialization process.

These 100 sequences were then used as input to DevRep and their predicted developability recorded.

After initialization, the sampling proceeds by first suggesting a Monte Carlo step and either accepting or rejecting the step according to the respective sampling criteria. In this proposed step, each of k residues in each sequence are mutated with a uniform probability from a pool of allowed amino acids. At the start of sampling, k=16 (all) positions in the sequence are mutated and changes in stringency (k) are made based on an arbitrary acceptance criterion.

For Nested Sampling (NS), at the beginning of each iteration, the lowest predicted developability is set as the threshold below which no sequences will be accepted. During each iteration we perform a short random walk during which each walker’s (mutated sequence) developability is predicted and compared to the threshold developability. We also assign an auxiliary random value uniformly sampled between 0 and 1 to each walker at each step. The developability of the proposed sequence is compared against the threshold with relative and absolute tolerance parameters of $10^{-5}$ and $10^{-8}$, respectively. If the developability is higher, the step is accepted, else it is rejected. In the case of a tie, we accept only if the auxiliary random number for the walker is larger than that of the threshold (in other words we sort sequences by developability first and then by this auxiliary random number). For each NS iteration we perform a random walk consisting of 5 such steps, after which we update the threshold and perform another NS iteration (and so on). Sampling terminates when the 100 mutating walkers collapse to yield a single unique most developable sequence.

For Simulated Annealing (SA), the walker's proposed step is accepted or rejected based on a Metropolis criterion with probability min(1, exp(-(new mutant developability – original developability)/Temperature)). Every walker is evolved at the same temperature for 550 steps after which the temperature is decreased. The temperature ramp is evenly spaced on a logarithmic scale from 10 to $10^{-3}$. The decrease of temperature is equivalent to increasing evolutionary pressure. Sampling terminates when the 100 mutating walkers have reached the end of the annealing process (*viz.*, the end of the decreasing logarithmically-spaced temperature ramp).

During each NS iteration and SA annealing step we record the average acceptance of the Monte Carlo steps. If the average acceptance is more than 30%, the number of k mutations in each step is increased by 1. If the average acceptance is below 20% then k is decreased by 1.

#### 2.4 *Analysis of Nested Sampling data*

The output of nested sampling is a list of threshold developabilities (and sequences) from which we can compute the density of states, $g(Y)$, and from it all thermodynamic observables such as the average yield, $\langle Y\rangle_\beta = \sum_Y Y g(Y)\, e^{-\beta Y} \;/\; \sum_Y g(Y)\, e^{-\beta Y}$, heat capacity $C(\beta) = \beta^2(\langle Y^2\rangle - \langle Y\rangle^2)$, entropy $S(\beta) = \beta(\langle Y\rangle_\beta - F(\beta))$ and free energy $F(\beta) = -\beta^{-1}\, ln(\sum_Y g(Y)\, e^{-\beta Y})$ where we have taken $k_B = 1$ everywhere. To generate the disconnectivity graphs, we first embed the recorded threshold sequences. Each of the embedded threshold sequences is connected to its K-nearest neighbors with (strictly) lower predicted yield (we choose K=2 and identify neighbors using the Euclidean distance from sklearn.Neighbors). If there are more than K sequences within the radius defined by the K-nearest neighbors (this happens if there are degeneracies, i.e. repeated identical threshold sequences), we connect the query sequence to all of these degenerate lower yield sequences. This procedure is repeated over all threshold sequences every 50 (for efficiency), so that the output is a graph in which each threshold sequence is a node connected to K (or more in case of degeneracies) lower developability neighbors. Then, we successively remove nodes and edges by going through the ordered list of thresholds yields (from low to high yields). When the graph splits into disconnected graphs, each subgraph is identified as a new basin (with relative sizes determined by the number of nodes). Knowledge of these subgraphs, with the predicted yields and the corresponding density of states of the nodes (threshold sequences) are used to plot the disconnectivity graph shown in Figure 8e [2].

### 3. *In vitro* Analysis

#### 3.1 *Gp2 Test Variant Selection*

From each Nested Sampling/Simulated Annealing run using OneHot, Unirep, and DevRep, all saved unique sequences were loaded. The sequences were embedded and reduced into their UMAP coordinates and then clustered using hbdscan. 100 sequences from each model/embedding were ordered, evenly split across the resulting clusters: For every cluster, we took the best 100/(number clusters) sequences to result in 100 diverse high developable variants total, excluding outliers. Additionally, we recorded 100 additional variants each for both from UniRep and DevRep; in this procedure, the top cluster for each embedding technique was further broken into smaller clusters and again sampled across the top of each group.

Therefore, for each strain, 100 (diverse) variants were recorded for each of the three embeddings (DevRep, UniRep, and OneHot) (300 variants total); 100 randomized-paratope variants; and 100 (developability-centric) variants were recorded for each of the DevRep and UniRep embeddings (200 variants total). This procedure thus resulted in an experimental variant pool of 600 Gp2 variants per each of the two *E. coli* strains.

#### 3.2 *Gp2 Test Variant Codon Design*

The amino acid sequence of each paratope was independently codon optimized for production in *E.coli* using IDT's Codon Optimization Tool. We then stitched the paratopes together with the codon-constant conserved regions, and a flanking sequence for PCR amplification.

#### 3.3 *Gp2 Oligopool*

Oligopools (Twist Bioscience) were designed to transform *E. coli* strains with plasmids encoding our 600 Gp2 variants as previously described[1].

Note that while our oligopool attempted to produce all 600 Gp2 variants for each of our two strains, only 280/600 and 269/600 variants were characterized for the $I^q$ and SH strains, respectively. We hypothesize this discrepancy in attempted versus successfully measured variants as arising from the stochasticity inherent in sampling from a large combinatorial space of potential oligonucleotides in our pool.

#### 3.4 *Gp2 Production for Dot Blot*

Dot blot was performed as a modified western blot with higher throughput. Six deep 96 well plates were prepared with 500 μL LB + Kan, wells were inoculated from previously prepared frozen stocks, and they were incubated overnight (SHuffle T7 Express LysY cells were grown at 30 °C and T7 Express LysY/$I^q$ cells [New England Biolabs] at 37 °C for the entire production). The next day, 25 μL per well of those cultures was added to freshly prepared deep well plates with 1 mL LB + Kan and grown for 90 min. Induction followed by the addition of 0.5 mM IPTG and incubation for 2 hrs ($I^q$) or 4 hrs (SH). Cells were then centrifuged (3000 x g, 5 min), supernatant was removed and pellet frozen at -80 °C. Pellet was thawed by the addition of 100 μL lysis buffer (20 mM sodium chloride, 2 mM magnesium chloride, 25 mM imidazole, 5 mM HEPES, 1 protease inhibitor pellet [Thermo Fisher Scientific], 2 μL benzonase nuclease [Millipore Sigma], 5 mg lysozyme [Millipore Sigma] in 50 mL $H_2O$ set to pH 8). Cells were lysed by incubation at 37 °C for 1 hr, then centrifuged for 5 min at 3000 x g. Supernatant from each well was added to a fresh 96 well PCR plate with 25 μL denaturing buffer (1 g/L SDS, 500 mM sodium chloride, 20 mM imidazole, 20 mM HEPES, pH 7.4). Plates were stored at 4 °C overnight.

#### 3.5 *Dot Blot*

Plates were removed from fridge and reheated for 5 min at 70 °C. PVDF membrane (0.2 μm pore, BioRad) for blotting were cut and placed into a box. The membrane was soaked in 50 mL methanol for 30 sec, then 50 mL d$H_2O$ for 2 mins. The membrane was then equilibrated via addition of 50 mL TBST (0.05% vol/vol Tween 20 in tris-buffered saline). The membrane was placed onto a TBST-soaked filter paper and dried with a Kimwipe. Using a multichannel pipet, 2 μL/well of protein sample was added to the membrane. After allowing the dots to fully absorb, the membrane was transferred to a fresh filter paper and allowed to dry in the fume hood for 30 minutes. The membrane was placed back into the box with 50 mL blocking solution (5% wt/vol nonfat dry milk in TBST) and rocked overnight at 4 °C. The membrane was labeled with 50 mL 0.2 μg/mL anti His6–horseradish peroxidase (Abcam) in blocking solution for 30 min at room temperature. Excess antibody was washed via 3x 50 mL TBST washes (room temperature for 10 minutes). The membrane was then soaked in 25 mL SuperSignal West Pico PLUS Chemiluminescent Substrate (ThermoFisher) for 5 min. The membrane was then placed inside a transparency and imaged via 5-30 sec exposure on a ChemiDoc MP Imaging System (BioRad).
